# Diversity and evolution of surface polysaccharide synthesis loci in Enterobacteriales

**DOI:** 10.1101/709832

**Authors:** Kathryn E. Holt, Florent Lassalle, Kelly L. Wyres, Ryan Wick, Rafal J. Mostowy

## Abstract

Bacterial capsules and lipopolysaccharides are diverse surface polysaccharides (SPs) that serve as the frontline for interactions with the outside world. While SPs can evolve rapidly, their diversity and evolutionary dynamics across different taxonomic scales has not been investigated in detail. Here, we focused on the bacterial order Enterobacteriales (including the medically-relevant Enterobacteriaceae), to carry out comparative genomics of two SP locus synthesis regions, *cps* and *kps*, using 27,334 genomes from 45 genera. We identified high-quality *cps* loci in 22 genera and *kps* in 11 genera, around 4% of which were detected in multiple species. We found SP loci to be highly dynamic genetic entities: their evolution was driven by high rates of horizontal gene transfer (HGT), both of whole loci and component genes, and relaxed purifying selection, yielding large repertoires of SP diversity. In spite of that, we found the presence of (near-)identical locus structures in distant taxonomic backgrounds that could not be explained by recent exchange, pointing to long-term selective preservation of locus structures in some populations. Our results reveal differences in evolutionary dynamics driving SP diversity within different bacterial species, with lineages of *Escherichia coli*, *Enterobacter hormachei* and *Klebsiella aerogenes* most likely to share SP loci via recent exchange; and lineages of *Salmonella enterica*, *Citrobacter sakazakii* and *Serratia marcescens* most likely to share SP loci via other mechanisms such as long-term preservation. Overall, the evolution of SP loci in Enterobacteriales is driven by a range of evolutionary forces and their dynamics and relative importance varies between different species.

## Introduction

Polysaccharide capsules and lipopolysaccharides (LPS) with an O-antigen, here broadly called surface polysaccharides (SPs), are the most diverse bacterial cell surface structures. They play a number of important biological roles pertaining to bacterial survival, including prevention from desiccation ^1, 2^, aiding transmission and colonisation ^3, 4, 5,6^, evading immune responses ^7,8,9,10^ or bacteriophage attack ^11,12,13,14^, interaction with other microorganisms ^15,16,17,18^, and many others ^19^. As SPs have been found and described in most studied phyla across the bacterial kingdom, their importance is recognised across many fields of biology, including ecology, medicine, biotechnology and public health^19,20,21,22^.

The potential for structural hyper-diversity of SPs stems from a heterogeneous ability of forming chemical linkage between various sugars into polysaccharide chains. This property was revealed in early work on bacterial carbohydrates ^23^, and understood in much more detail following the emergence of genetics as a field^24^. Epidemiological characterisation of bacterial serotypes ^25,26,27^ played an important role in building accurate models of the SP genotype-phenotype map in multiple bacterial species ^28,29,30^, hence enabling *in silico* serotyping approaches ^31, 32^. Today, we understand that the potential to generate novel SP diversity is genetically optimised in bacteria: co-location of genes that encode sugar-specific enzymes facilitates allele and gene transfer via homologous recombination between different bacteria, thus enabling antigenic diversification ^33^.

Nevertheless, our understanding of SP evolution remains far from complete. SP genetics has predominantly been studied in a small number of medically-relevant bacterial species, with little attention paid to comparative evolutionary dynamics or SP sharing between species. For example, it remains unclear how SP biosynthesis loci have been evolving in the context of different bacterial population backgrounds (whether defined by ecology or phylogeny), whether the long-term impact of horizontal gene transfer (HGT) on SP loci is the same in different taxonomic groups, and how selection on SP loci shapes bacterial population structure. These questions are important to answer since capsules and LPS directly interact with the immune systems of humans and other mammalian hosts, and are targets of current and future medical interventions, including glycoconjugate vaccines ^34^ and antibody^35^ or phage-based therapies ^36^. A better understanding of the diversity and evolutionary dynamics of bacterial SPs and of their role in bacterial adaptation to novel ecological niches could therefore have large public health impacts in terms of infectious disease management, for instance through the assessment of which serotypes to include in a vaccine to minimise the risk of disease reemergence ^37^.

Here we present an analysis of SP locus diversity and evolution in a bacterial order of medical importance – Enterobacteriales, which includes the well-known Enterobacteriaceae family (including *Escherichia*, *Salmonella*, *Klebsiella*, *Enterobacter*) and the related families Erwinaceae (including *Erwinia*), Yersinaceae (including *Yersinia* and *Serratia*) and others which were recently removed from the Enterobacteriaceae family definition ^38^. This group constitutes a good system to study the evolutionary genetics of polysaccharide capsules for two main reasons. First, many Enterobacteriales species have a closely related capsule genetic architecture ^39^, and instances of SP gene sharing between different genera in this order have been noted^40, 41^. Second, in recent years there has been a rapid growth of public genome collections of Enterobacteriales, largely due to the increasing threat of antimicrobial resistance in the Enterobacteriaceae^42^. Therefore, public repositories potentially include many isolates of Enterobacteriales species with previously uncharacterised SP genetics. Nevertheless, detailed analyses of polysaccharide genetic variation have been confined to a small number of species ^39, 43,44,45,46,47^. Here we used a large collection of 27,334 genomes obtained from NCBI RefSeq covering 45 genera of Enterobacteriales to carry out order-wide comparative genomics of two well-characterised SP locus regions, here referred to as *cps* and *kps*, which are involved in the biosynthesis of both capsules and O-antigens in *Escherichia coli* ^48^ and in other species of the Enterobacteriaceae family ^44, 49, 50^ (see Figure 1A for well-studied examples of *cps* and *kps*). We explore evolutionary dynamics and horizontal transfer within and between species and genera of Enterobacteriales, yielding the largest-to-date systematic analysis of the SP locus genetics in any bacterial family or order.

**Figure 1.**
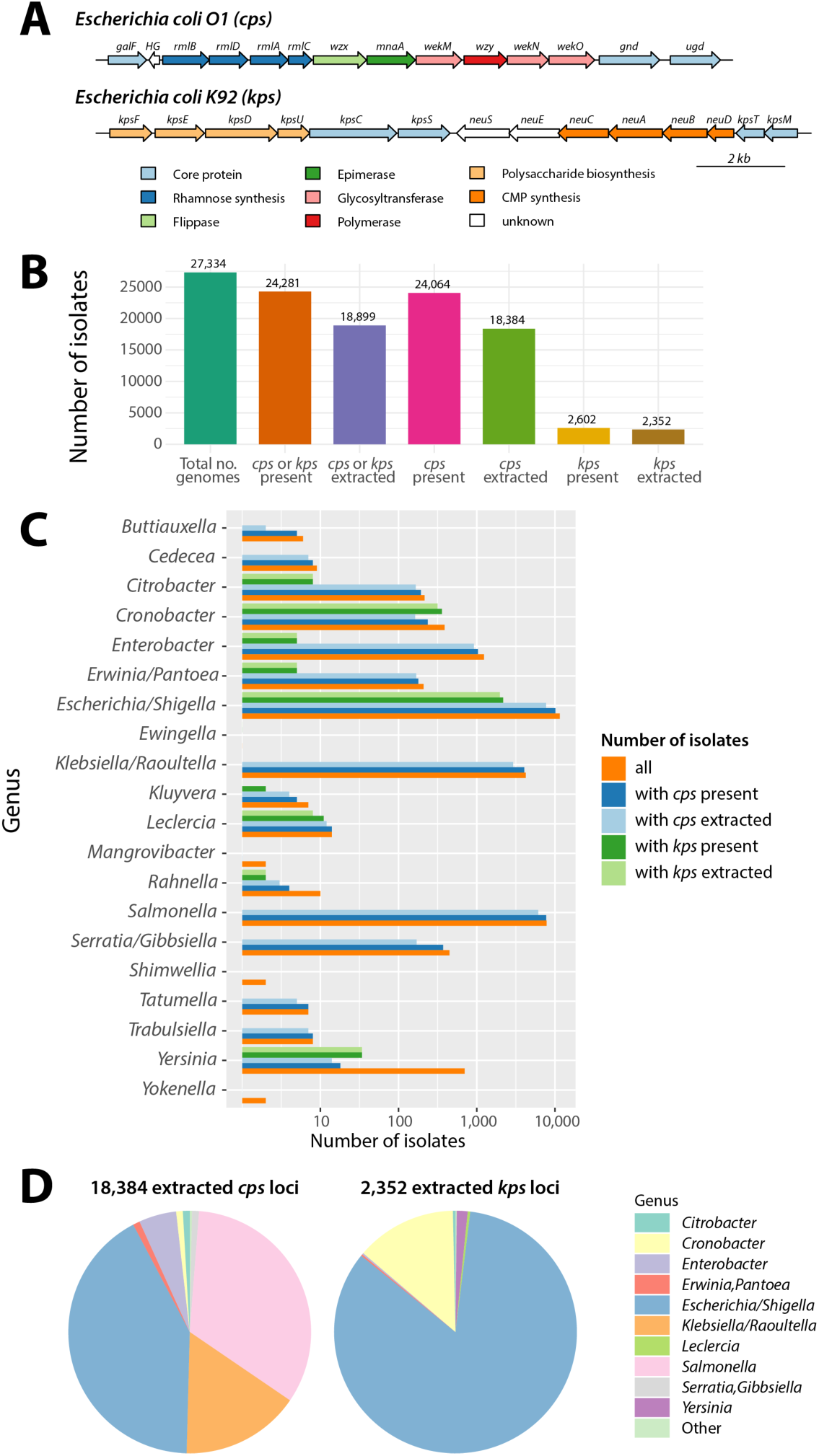
Details of SP locus extraction. (A) Overall genomic architecture of the *cps* (top) and *kps* (bottom) locus regions, based on well-studied examples of *Escherichia coli* O1 serotype and *Escherichia coli* K92 serotype. Genes are coloured according to the main functional classes with core genes used as locus markers shown in light blue (see Methods). (B) Locus presence/extraction statistics in the studied dataset. From the left, the bars show the total number of QCed genomes (green), number of isolates with either locus present (orange) and extracted (purple), number of isolates with the *cps* locus region present (magenta) and extracted (olive green), number of isolates with the *kps* locus region present (yellow) and extracted (brown). (C) Bar plot shows the number of isolates within each genus-group in the following categories: total number of isolates (orange), number of isolates where the *cps* locus region was present (dark blue), number of isolates where the *cps* locus region was present and of high enough quality to extract (light blue), number of isolates where the *kps* locus region was present (dark green), number of isolates where the *kps* locus region was present and of high enough quality to extract (light green). Only genus-groups in which at least a single *cps* or *kps* locus was detected are shown. (D) Pie charts show the relative frequency of loci extracted from different genera in the *cps* locus region (left) and the *kps* locus region (right). The frequency for genus X was calculated as the number of isolates of genus X with the *cps* (*kps*) locus region extracted to the total number of isolates with the *cps* (*kps*) locus region extracted. Only genera with frequency greater than 10*^-^*^3^ are shown by their name, otherwise they were pooled into the ‘Other’ category.

## Results

### Diversity and distribution of SP loci

Screening 27,334 assemblies of Enterobacteriales from 45 genera (see Methods and Supplementary Figure S1 illustrating the workflow) identified a high quality *cps* locus in 18,384 genomes from 22 genera (counting *Escherichia* and *Shigella* as a single genus), and a high quality *kps* locus in 2,352 genomes from 11 genera; the remaining genomes either contained poorly assembled locus sequences or were missing altogether (see Figure 1). The supplementary figure S2 shows the frequency of *cps* and *kps* in different genus-groups, demonstrating that the vast majority (92%) of genomes with *kps* also carried *cps*. The number of unique SP loci (i.e., comprising unique combinations of protein coding sequences, CDS) detected per genus was strongly predicted by sample size (number of genomes analysed per genus), but not the nucleotide diversity captured by that sample (see Supplementary Text S1 and Supplementary Figure S3), consistent with recent observations in *Klebsiella pneumoniae* ^51^. Consequently, we predict large reservoirs of unobserved SP diversity to be discovered and characterised as more genomes are sequenced. Overall, most locus types (groups of highly similar SP loci, defined by clustering with gene content Jaccard distance 6 0.1; see Methods) and SP locus gene families (LGFs, homology groups identified in SP loci and clustered at 50% amino acid identity; see Methods) were species specific: in the *cps* region, 90% of locus types and 61% of locus gene families were found in a single species, while in the *kps* region it was 93% and 78%, respectively. However, we also found the same locus types in multiple species and genera (*cps*: 9.8% in multiple species and 3.3% in multiple genera; *kps*: 7.1% in multiple species and 0.89% in multiple genera), suggesting that the evolutionary history of SP loci involves horizontal transfer across genus boundaries (see Supplementary Figure S4).

To further explore the diversity and population structure of SP loci, we constructed locus-sequence similarity networks for the *cps* and *kps* loci extracted from the genome set (Fig. 2A). To avoid redundancy, we considered a set of loci consisting of single representatives each SP locus structure (i.e., each detected combination of LGFs) from each species (*N* = 2,654 *cps*, *N* = 332 *kps*; see Methods). We then defined a locus similarity network for a given Jaccard distance threshold, *J*, where nodes are representative loci and edges link all loci whose gene content similarity is equal or greater than *S* = 1 *- J*. For the networks shown in Figure 2A, the similarity threshold was chosen to maximise the clustering coefficient (see Fig. 2B), which measures the degree to which nodes in a graph tend to cluster together. The large majority of communities (i.e., clusters of linked nodes with similar gene content) in each network comprise SP loci from the same genus; however many genera had SP loci that were divided into multiple unconnected communities. Many between-genus links were also observed (14% in both the *cps* and the *kps* network), which did not disappear even for *S* = 1 (5% for *cps* and 2% for *kps*; see Fig. 2B and Methods), indicating the presence of SP loci with identical gene content in distinct bacterial genera.

**Figure 2.**
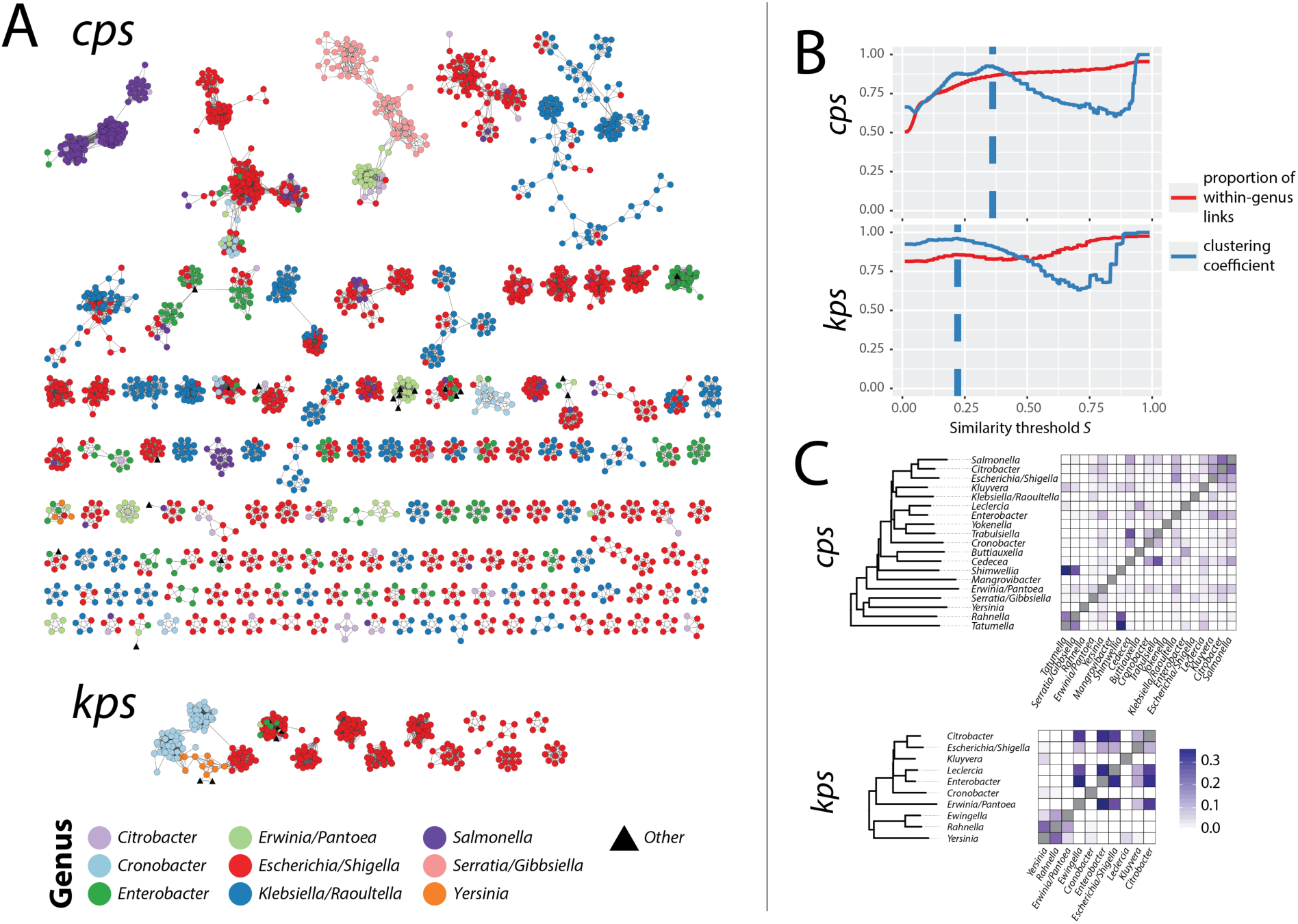
Population structure of SP loci is weakly correlated with the order population structure. (A) SP locus sequence similarity networks (SSNs). Each node corresponds to a unique locus genetic structure per species (*n* = 2654 nodes for *cps*, *n* = 332 nodes for *kps*; only connected components consisting of at least 5 nodes are shown). Colours correspond to bacterial genera in which loci were found (9 most common genera and ‘Other’). Edges of the network link all nodes *a* and *b* for which the similarity threshold *S*(*a, b*) = 1 *J*(*a, b*) maximises the clustering coefficient (dashed lines in B), namely *S*(*a, b*) > 0.362 for *cps* and *S*(*a, b*) > 0.222 for *kps*. Connected components in this graph define locus families, while connected components of near-identical locus structures (*S* > 0.9) define locus types. (B) Effect of the similarity threshold on SP locus clustering. (C) SP locus gene sharing between genera. Dendrograms are neighbour-joining trees based on whole-genome genetic distance between representative (non-redundant) members of each genus; heatmaps show the proportion of SP locus gene families (LGFs, defined at 50% clustering threshold) that are shared between each pair of genera. In this figure, the most prevalent LGFs (present in > 20% of all isolates) were removed from the calculation of the locus similarity to amplify the signal from the low-to-middle frequency protein families.

We observed a large heterogeneity in LGF sharing between genera: some genus pairs shared large numbers of SP genes while others shared none (see Fig. 2C). These LGF sharing patterns however showed no clear association with taxonomic relationships between the genera in question, and indeed some of the strongest LGF sharing occurred between members of different families (e.g., *Shimwellia* (Enterobacteriaceae), *Tatumella* (Erwinaceae) and *Rahnella* (Yersinaceae) shared very similar *cps* gene complements; Erwinia (Erwinaceae) shared very similar *kps* gene complements with members of the Enterobacteriaceae family; see Fig. 2C). A similar analysis at the species level (see Supplementary Figure S5) showed that these patterns were not driven by a small number of species; there were many pairs of distant species that shared low-to-middle frequency LGFs. Furthermore, we found weak evidence of correlation between the genome-wide genetic distance between genera (based on average nucleotide identity, ANI across non-redundant representative genomes) and the Jaccard similarity *S* between sets of LGFs detected in those genera: borderline significantly positive correlation for the *cps* region (Spearman rank correlation; *ρ* = 0.16, *p* = 0.031) and insignificant correlation for the *kps* region (Spearman rank correlation; *ρ* = 0.20, *p* = 0.20); see also Supplementary Figure S6. We observed an even weaker evidence for a locus and genome population structure at the species level (see Supplementary Figure S5): a weak correlation for the *cps* region (Spearman rank correlation; *ρ* = 0.038, *p* = 0.0071) and no significant correlation for the *kps* region (Spearman rank correlation; *ρ* = 0.031, *p* = 0.56).

### Mechanisms of locus type sharing between divergent strains

To better understand the dynamics of locus type sharing between divergent strains, we calculated a probability of sharing the same locus type between representative isolates from different *L*_0.004_ lineages as a function of genome distance. (Representative isolates were chosen as unique combinations of ribosomal sequence type and unique locus structure, *L*_0.004_ lineages were defined as complete-linkage clusters distinct at the 0.4% genome distance threshold, and genome distance was calculated based on estimated ANI values; see Methods.) Results are shown in Figure 3A. As expected, the probability of locus type sharing decreased with genome distance for both *cps* and *kps* (with the exception of *kps* around 8% genome divergence, driven by a high proportion of shared locus types between different species of *Cronobacter*), followed by a long tail expected from Fig. 2. We hypothesised that many instances of locus sharing may be driven by recent horizontal exchanges of the whole (or nearly-whole) locus via HGT. To address this, we extracted all pairwise genome combinations belonging to different *L*_0.004_ lineages that shared a locus type, and compared genome vs SP locus genetic distance between each pair (Figure 3B). While whole-genome and SP locus distances were strongly correlated (*cps*: *R*^2^ = 0.90, *p <* 10*^-^*^16^; *kps*: *R*^2^ = 0.71, *p <* 10*^-^*^16^), for many pairs the SP locus genetic distance was smaller than the whole-genome genetic distance, as would be expected from HGT-driven locus exchanges (Figure 3B). To examine to what extent locus type sharing between *L*_0.004_ can be explained by recent horizontal exchanges, we calculated, for each locus-type-sharing pair, where the SP locus genes ranked within the distribution of pairwise distances between protein sequences for all CDS shared by that pair (see Methods). We reasoned that, if the SP locus has been exchanged (directly or indirectly) between two diverged lineages via recombination, the SP locus genes should rank amongst the most genetically similar CDS for that pair of lineages (see also Supplementary Figure S7). (Note that by ‘recent exchange’ we mean an HGT event that occurred relatively recently in comparison to the evolutionary distance between two lineages considered. Furthermore, across the Enterobacteriales population, SP locus genes are overall less conserved than other genes – see Supplementary Figure S8 – hence this approach is unlikely to result in false-positive calls of recombinational exchange.) We considered locus-type-sharing lineage pairs in which the SP loci rank in the top 5% most similar CDS for that pair as likely resulting from horizontal exchange of the SP locus between lineages relatively recently (since the divergence of the lineages) (blue); and those locus-type-sharing pairs in which the SP loci rank in the bottom 60% most similar CDS for that pair as evidence of absence of recent horizontal exchange (red); the remaining cases we considered unresolved (grey). Using such criteria, we found 705 pairwise instances of locus type sharing between species, and 218 between genera, that were attributable to recent exchange affecting the *cps* locus region; plus 59 and 2 cases, respectively, affecting the *kps* locus region. While our approach does not necessarily allow us to distinguish which of these instances are truly independent, their visualisation in the form of a network (see Figure 4) shows the extent to which SP locus diversity can and has spread via HGT across Enterobacteriales. Four examples of between-genus locus exchanges (three for *cps* and one for *kps*) are explicitly shown in Supplementary Figure S9. Instances of locus type sharing in the absence of recent exchange are shown in Supplementary Figure S10.

**Figure 3.**
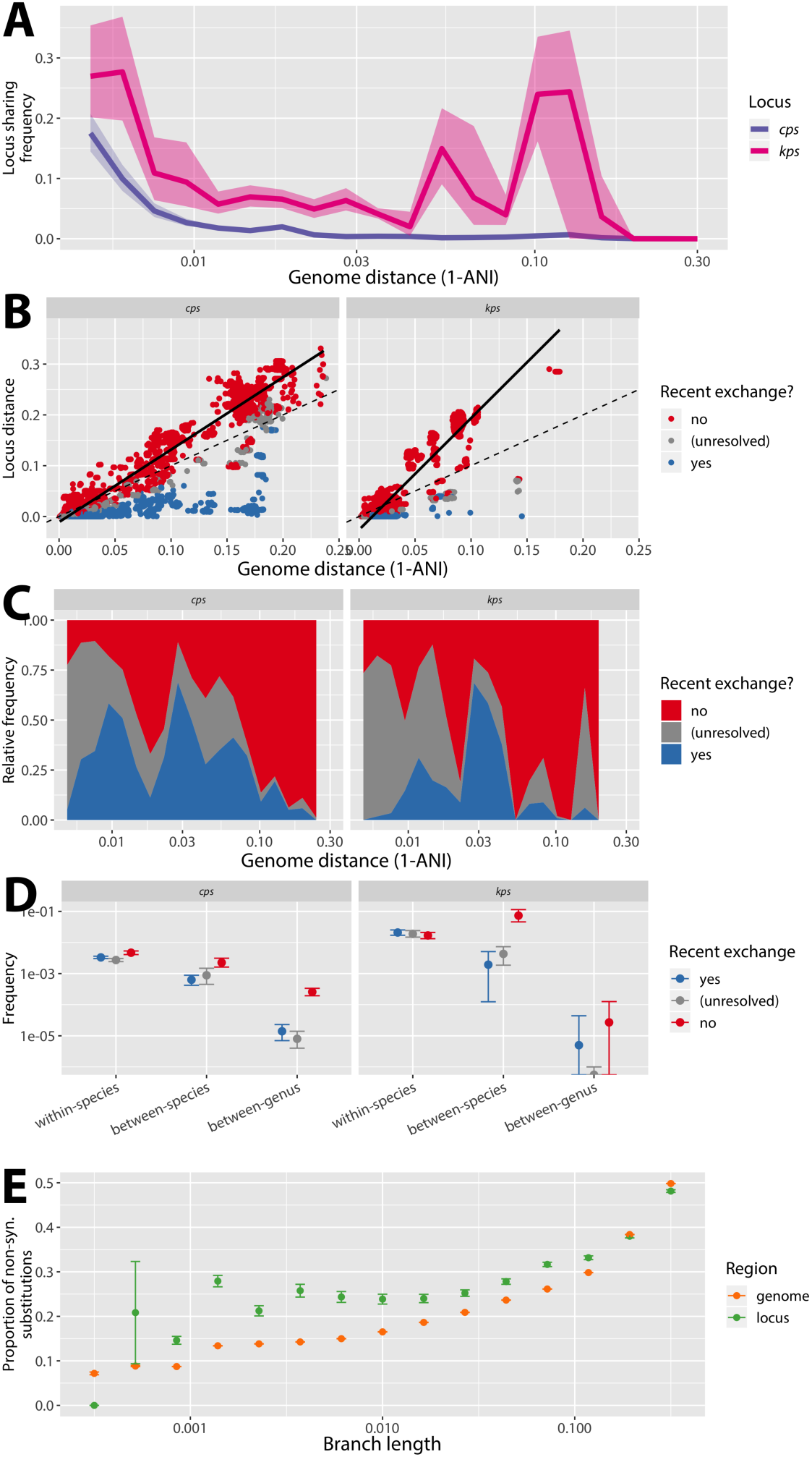
SP locus type sharing between distant members of the Enterobacteriales. (A) Estimated probability of sharing a locus type between representative isolates from different *L*0.004 lineages as a function of their genome distance based on average nucleotide identity (ANI). Curves were calculated by considering *n* = 20 ANI distance ranges; and for all isolate pairs with genomic distance within that range, calculating the proportion of all pairs that share the SP locus type. Shaded areas indicate the 95% confidence intervals obtained by bootstrap, based on random isolate sampling with replacement repeated 1,000 times; solid line shows median value of the bootstrap distribution. (B) Pairwise genetic distance between SP loci (*y*) and bacterial host genomes (*x*), using estimated ANI, for all isolates from different *L*0.004 lineages that share a locus type. Data points (strain pairs) were assigned to categories: evidence of recent locus exchange (blue), evidence of no recent locus exchange (red) or unresolved cases (grey), based on the relative distance of SP locus genes vs genome-wide CDS (see Methods). Dashed line is *y* = *x*, solid line is the linear fit to all red points (*cps*: *R*^2^ = 0.97, *p <* 10*^-^*^16^; *kps*: *R*^2^ = 0.87, *p <* 10*^-^*^16^). (C) Relative contribution of recent exchange presence (blue), recent exchange absence (red) and unresolved case (grey) to locus type sharing as a function of genome distance. (D) Estimated probability of locus type sharing via one of the three categories by taxonomic level. Values were calculated as in panel C-D, but stratified by taxonomic level rather than evolutionary distance range. (E) Proportion of total substitutions per gene tree branch that are non-synonymous, for SP locus genes (green) and other genome-wide CDS (orange); stratified by branch length ranges (x-axis). Bars indicate 95% confidence intervals (using one sample test of proportions).

**Figure 4.**
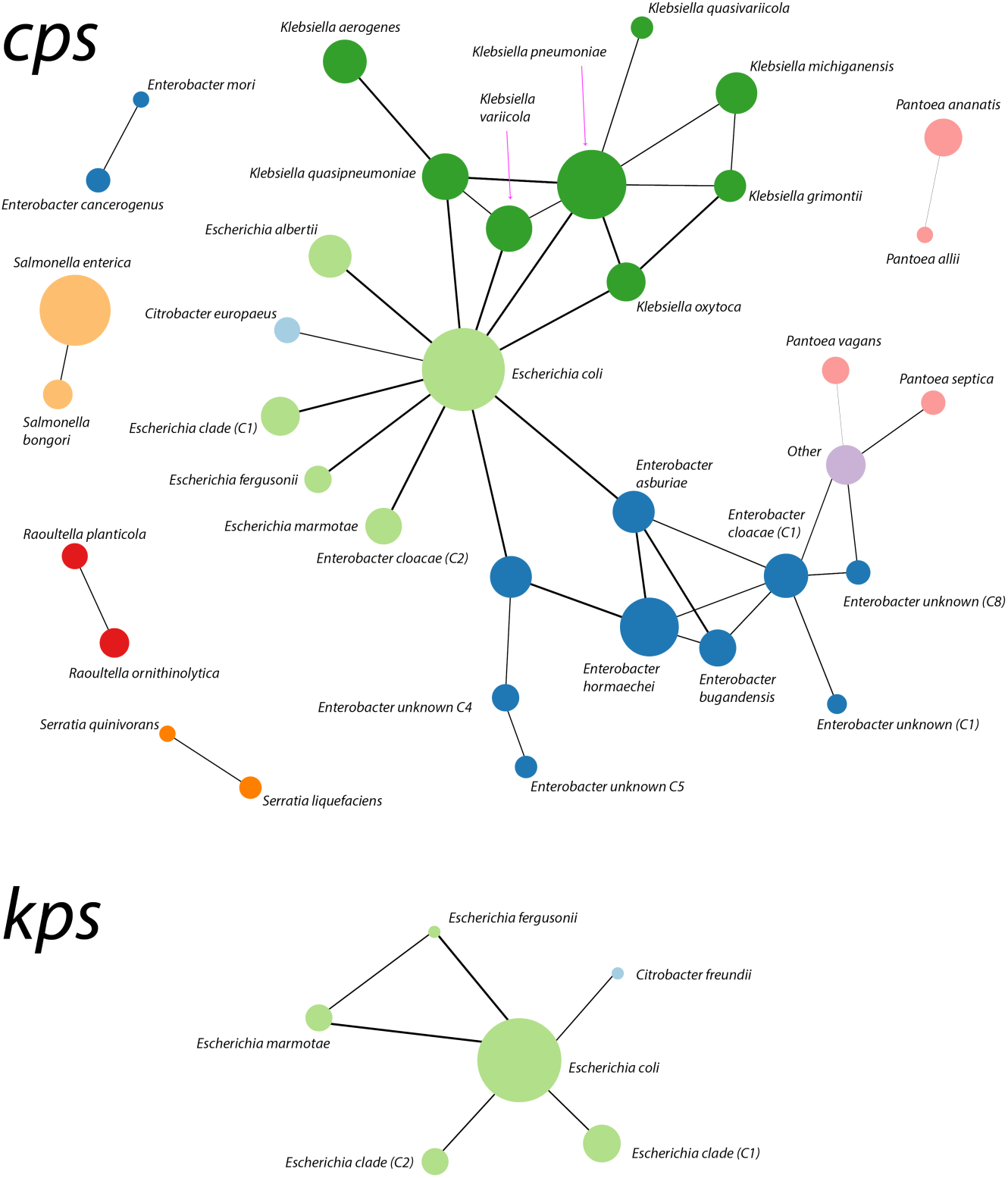
Network of SP locus type sharing via recombinational exchange. Recombinational exchange of SP locus was defined as explained in the main text (blue points in Figure 3). Nodes represent bacterial species-groups, and their size is scaled according to the number of isolates in the representative dataset with the corresponding locus region (*cps* or *kps*). Edges represent cases of detected locus exchanges, and their thickness is scaled depending on the number of detected cases of sharing. Using such definitions, we found 705 cases of shared *cps* loci between different enteric species and 218 cases between different genera (699 and 218 when excluding ‘Other’ category, respectively). For the *kps* locus, we found 19 between-species sharing cases and 2 between-genus sharing cases (same when excluding ‘Other’ category).

Figure 3C shows the relative contribution of the three categories to locus type sharing as a function of genome distance. We found that recent locus exchange can explain as much as 50% of locus type sharing cases for genomic distance of up to around 10%, and even some cases for distance *>* 10%. The estimated probability of locus type sharing via recent exchange was greatest within species, but greater than zero even between species and between genera (again except for *Cronobacter*-driven between-species exchanges in the *kps* locus; see Figure 3D). For closely related genomes, the majority of locus type sharing fell into the grey category, highlighting the limited statistical power to detect recent exchange between recently diverged backgrounds (Fig. 3C). Conversely, locus type sharing between distant lineages (*>* 10%) was predominantly not attributable to recent exchange (Fig. 3B and 3C). Such occurrences could arise through vertical inheritance of SP loci over long evolutionary periods, or by independent acquisition of divergent forms of the same locus type by different lineages (see Supplementary Figure S11 for an illustration of these scenarios). Whatever the mechanism, presence of the same locus types in highly divergent lineages is suggestive of a form of stabilizing selection that preserves the structure of SP loci over time, resisting genetic change in spite of high rates of HGT. We also investigated two factors as potential predictors of locus type exchange via HGT: population frequency and locus gene content (see Supplementary Text S2 and Supplementary Figures S12 and S13) but found no evidence of a straightforward relationship, suggesting that locus exchanges between bacterial lineages are driven by a complex interplay of selective and ecological forces interacting with the microbial population, rather than e.g. universal selection for spread of loci associated with a particular type of sugar.

For instances of locus sharing where we could reject the hypothesis of recent exchange, SP locus gene distance generally exceeded genome distance (i.e., most red points in Fig. 3B lie above the dotted line *y* = *x*). The slopes of the regression lines for these points were significantly greater than 1 (black lines in Fig. 3B; *cps*: *ff* = 1.426, 95% CI [1.423,1.428]; *kps*: *ff* = 2.198, 95% CI [2.179,2.216]). The SP locus genes also had more non-synonymous substitutions than other CDS in the Enterobacteriales genomes (45.0% of all substitutions in SP locus genes were non-synonymous compared with 26.3% for other CDS, *p <* 10*^-^*^15^), suggesting comparatively relaxed purifying selection at the SP loci. These observations could reflect an overall accelerated evolutionary rate in SP locus genes, or the acquisition of highly divergent gene copies by HGT. To address these biases, we calculated the proportions of non-synonymous substitutions on the branches of each gene tree (thus accounting for the effect of potentially distant origins of alleles), and binned these values by branch lengths across all HG trees (Fig. 3E). Across almost all branch length ranges, the SP locus genes had a higher proportion of non-synonymous substitutions than other CDS, supporting weaker purifying selection at SP locus genes irrespective of differences in overall evolutionary rate.

### SP locus evolution within species

Next we examined patterns of SP locus type sharing between lineages of the same species. Figure 5A shows the probability of the same locus type being present in two genomes (i.e., Hunter-Gaston diversity index^52^) whose distance is below a given threshold, across a range of thresholds from 10*^-^*^4^ to 0.3. For closely related genomes the probability was close to 1, but dropped off dramatically above 1% divergence, reaching different plateau probabilities in different species. For the two species with sufficient numbers of both *cps* and *kps* (*E. coli* and *C. sakazakii*), the two SP locus curves were similar within each species (Fig. 5A). The plateau probabilities could be influenced by sampling; in particular, over-sampling of some lineages, which is a common feature of public genome repositories, would lead to an over-estimate of the plateau probabilities. To mitigate this bias, we calculated for each species the probability that genomes from two different lineages share a locus type (Fig. 5B), using a resampling approach and two alternative thresholds to define lineages (*L*_0.004_ or *L*_0.01_). These measures were not significantly correlated with the sample size per species (labelled in Fig. 5B) (Spearman rank correlation for *L*_0.004_: *p* = 0.10, *L*_0.01_: *p* = 0.57), but confirmed significant differences in SP locus-sharing patterns between different species, with probability values ranging from 0.47% (95% CI: 0.44%, 0.51%) in *E. coli* to 56% (95% CI: 37%, 79%) in *K. aerogenes* using the *L*_0.004_ lineage definition.

**Figure 5.**
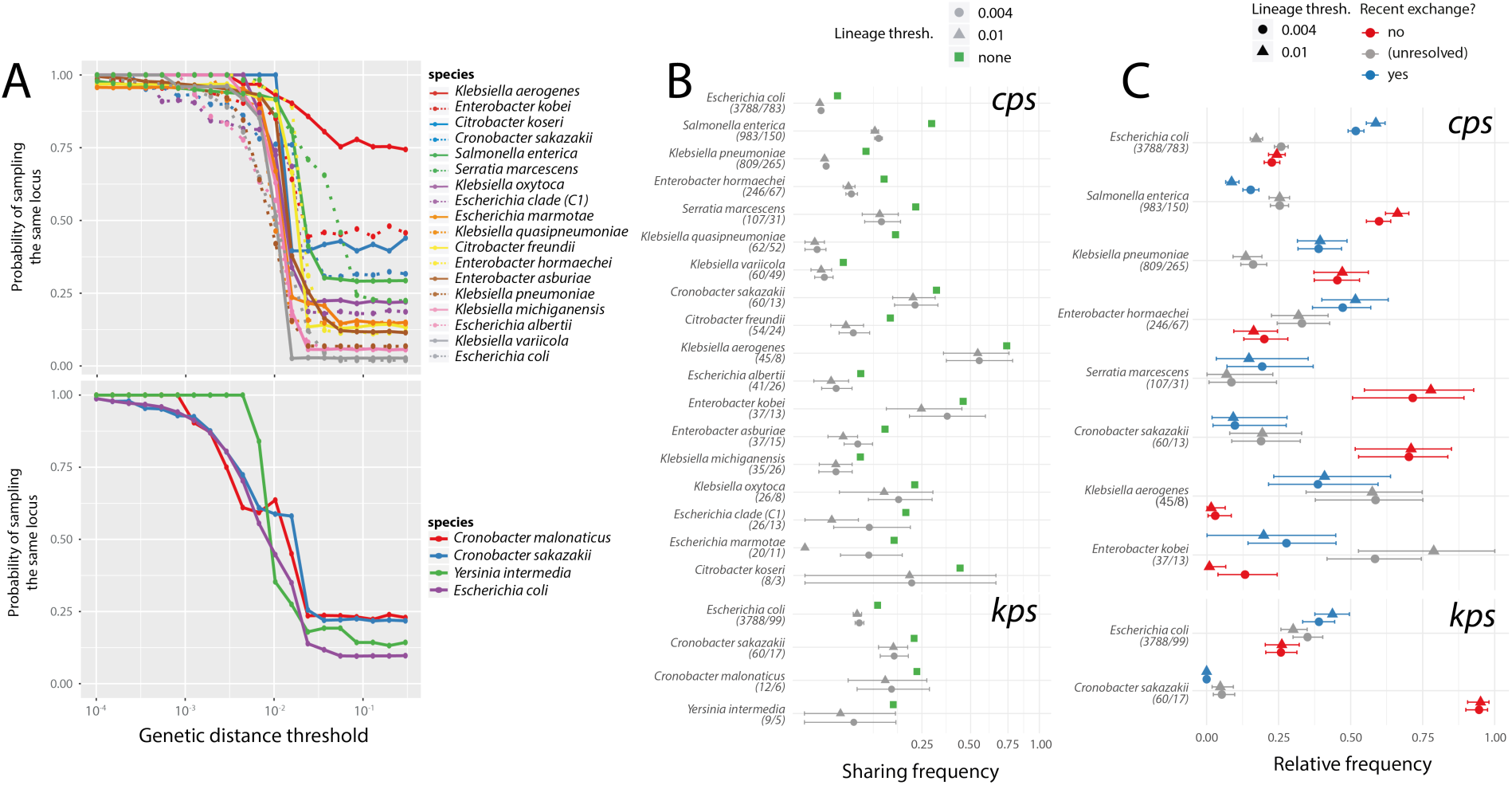
Intra-species patterns of SP locus exchange and maintenance vary between species. (A) Probability of locus type sharing in each species, as a function of the maximum genomic distance (only genome pairs of a given distance or less are compared). Results are shown for the 18 species in which our dataset includes at least 5 *L*0.01 lineages with mean >2 isolates per lineage. The values plotted are the median bootstrap values when resampling all isolates with replacement *n* = 100 times for a given genome distance. (B) Estimated SP locus type sharing probability between isolates belonging to different *L*0.004 and *L*0.01 lineages. Error bars denote 95% confidence intervals obtained by bootstrap (resampling isolates with replacement within the species *n* = 1000 times). Green squares show the plateau probability value from panel A, and species are sorted by sample size. Values in parentheses next to each species name are the number of genomes and the number of locus types for this species, respectively. (C) Relative contribution of the three categories from Figure 3 (blue: recent exchange, red: no recent exchange, grey: unresolved) to SP locus type sharing between lineages. Only species in which we found *>*100 pairs of lineages sharing SP locus types are displayed. Values shown represent median of the bootstrap distribution sampled *n* = 1000 times.

To assess the contribution of recent exchange to locus-type-sharing between lineages, we considered only those species in which locus-type-sharing was observed for at least 100 lineage pairs, and calculated the proportion of *L*_0.004_ or *L*_0.01_ lineage pairs in which locus type sharing was attributable to one of the three categories in Figure 3: recent exchange (blue), no recent exchange (red) or unclear (grey); see Fig. 5C. (See also Supplementary Figures S14-S17.) Using this approach, we found that in several species, including *K. aerogenes*, *E. hormaechei* and *E. kobei* and *E. coli*, recent locus exchange was the predominant mechanism of locus type sharing between different bacterial lineages. By contrast, in *S. enterica*, *S. marcescens* and *C. sakazakii*, locus type sharing could be rarely explained by recent locus exchange, hence pointing to vertical inheritance or independent acquisition as potential mechanisms. Finally, in *K. pneumoniae* both recent exchange and lack thereof contributed equally to locus type sharing. Species with the more diverse SP locus were more likely to share locus-type via recent exchange, however this correlation was not statistically significant (Supplementary Figure S18). These results point to important differences in the evolutionary dynamics of SP locus sharing between different species, and are consistent with previous studies of antigen variation in some individual species. For example, frequent exchange of K and O antigen loci has been observed within *E. coli* lineages ^53, 54^, whereas O antigen (*cps*) loci are maintained in *S. enterica* lineages to the point that lineage typing has been proposed as a substitute for serotyping ^44^. In *K. pneumoniae*, the *cps* K locus is a hotspot for recombination in many lineages, but in others is conserved for long evolutionary periods ^51^. Our results thus highlight that different populations of Enterobacteriales not only experience a variation of forces acting on the SP locus, but also that these forces vary in relative magnitude between populations.

### Evolutionary dynamics of individual SP locus genes

The *cps* and *kps* loci follow the typical structure of SP loci, comprising a core complement of genes required for SP expression, regulation and export (at the ends of the locus) and a variable complement of genes coding for assembly of oligo- and polysaccharides (in the middle of the locus). In addition to relocation of whole SP loci into new chromosomal backgrounds via HGT (explored in detail above; see Figure 4), evolution of SP loci involves diversification of the individual component genes through substitutions and recombination, as well as gain and loss of sugar transferase and synthesis genes to form new locus structures with different gene complements. To explore gain and loss of individual LGFs over short time scales, we inferred neighbour-joining trees for each *L*_0.004_ lineage, and used the GLOOME software to infer the ancestral patterns of SP locus gene gains and losses on each tree using maximum parsimony (see Methods). The results reveal many instances of gain or loss of small numbers of genes (1-3) within individual bacterial lineages (Fig 6A), demonstrating that evolution of locus genetic content can occur on relatively short epidemiological timescales. Interestingly, we did not observe any significant relationship between this measure of individual gene mobility (gain/loss) rate and the relative position of the genes in the locus (see Supplementary Figure S19). This is in contrast to previous observations that the central sugar processing genes are subject to higher frequencies of homologous recombination in *K. pneumoniae* ^46^ (and in the Gram-positive *S. pneumoniae* ^55^), although this could be due to insufficient power given the sample size and short timescales captured in our gene gain/loss analysis. We did however observe clusters of co-evolving LGFs (Figure 7), which were strongly associated with co-mobility in the population (i.e., sharing similar patterns of gain/loss; see Supplementary Figures S20 and S21), consistent with evolution of SP loci through reassortment of modular gene sets to create new locus types.

**Figure 6.**
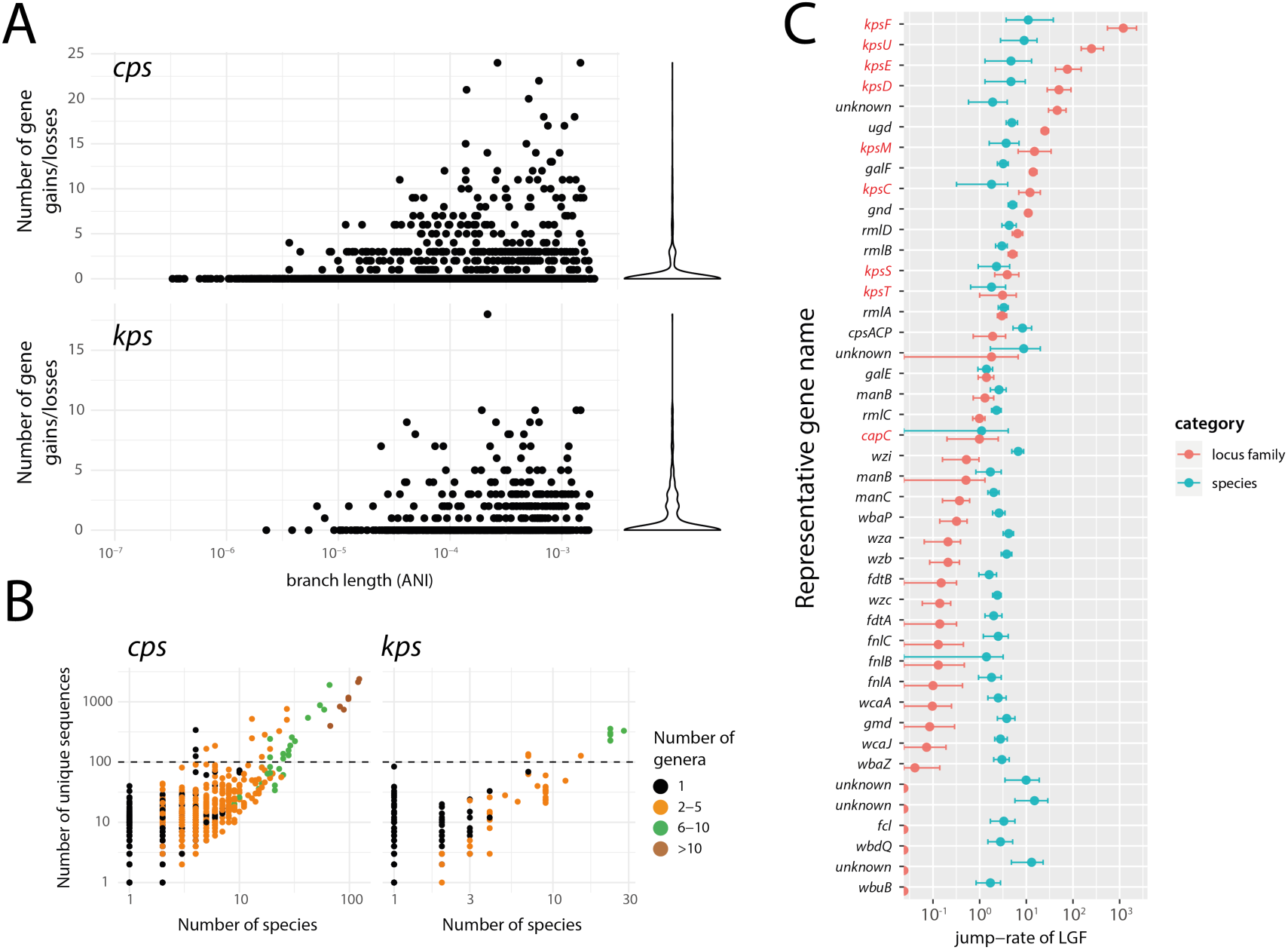
SP loci evolve via horizontal movement of individual locus gene families (LGFs). (A) Number of gene gains or losses per branch vs. branch lengths, across all *L*0.004 lineage trees (see Methods). The violin plot on the right-hand-side shows the marginal distribution of the number of gains/losses, dominated by zero (reflecting within-locus exchanges of small numbers of genes) and with a long tail (reflecting occasional whole-locus exchange events), detected within bacterial lineages. (B) Each point corresponds to one LGF with minimum two sequences (*n* = 3120 for *cps* and *n* = 283 for *kps*). Plot shows the number of species in which a given LGF was found vs the number of unique sequences in that LGF. (C) Species-jump-rates (blue) and locus-family-jump-rates (red) for LGFs with >100 sequences (above the dashed line in panel A). Bars indicate 95% confidence intervals obtained by bootstrapping (see Methods). LGFs are labelled with the most common gene name, *kps* LGFs are highlighted in red. Locus families were defined as connected components in Figure 2.

**Figure 7.**
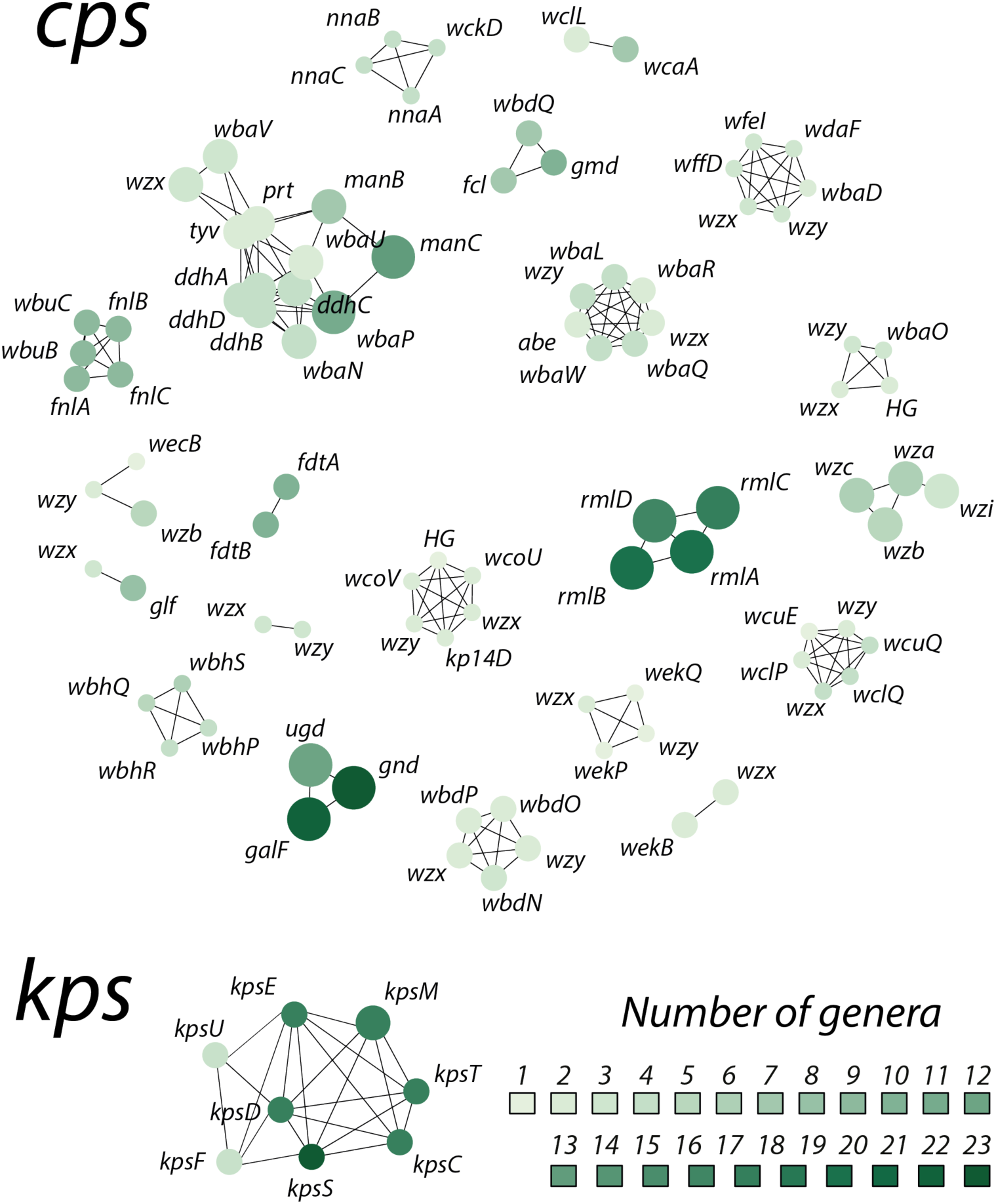
Co-evolution between locus gene families (LGFs). LGFs with pairwise co-evolutionary coefficient *CVij* > 0.5 and co-occurring in least 200 isolates are linked in the network. Nodes are scaled to indicate the number of isolates in which each LGF was found, coloured to represent the number of genera, and labelled with the most common gene name annotated for members of that LGF.

A large proportion of LGFs were found in diverse taxonomic backgrounds (Fig 6B): amongst all LGFs with at least two unique sequences, 62% (44%) of the *cps* (*kps*) LGFs were found in at least two species and 40% (24%) were found in at least two genera. We also found identical *cps* and *kps* LGF sequences in multiple species, indicating recent horizontal transfer of SP locus genes between diverse taxa. We hypothesised that different LGFs frequently move between different genetic backgrounds. To assess this, we quantified such moves as rates of jump (jump-rates) between species for individual LGFs using ancestral state reconstruction to model bacterial species as a discrete trait on each individual gene tree (for LGFs with *>*100 sequences), and measured the frequency of state transitions (i.e. species changes) per branch length (see Methods). The distribution of species-jump-rates for the most diverse LGFs (*>*100 sequences each; i.e. points above the dashed line in Fig. 6B) is shown as blue points in Figure 6C. The species-jump-rates were compared to locus-jump-rates, which were calculated as jump-rates between diverse locus families (connected components in Figure 2A), and are shown as red points in Fig. 6C. We found a much greater variance in locus-jump-rates than in species-jump-rates. The flanking *kpsFEDU* genes in the *kps* locus and *galF/gnd/ugd* genes in the *cps* locus were more likely to jump between locus families than between species. Amongst LGFs with the highest jump-rate, we found five genes with unknown function. Using hhsuite ^56^, we found that two of those genes had resemblance to distant enzyme protein sequences (phosphomannomutase and 5-formyltetrahydrofolate cyclo-ligase) in the UniProt database. These unknown genes could thus represent components of uncharacterised polysaccharide biosynthesis pathways; but may also potentially represent unknown mobile genetic elements (see Supplementary Table S2).

## Discussion

To our knowledge we present the first systematic analysis of SP locus evolution across Enterobacteriales, or indeed any bacterial order. Previous studies focusing on individual SP loci, or the distribution of SP loci within individual species groups, have occasionally reported instances of individual loci transferring across species or genus bounds. However this study provides a unique view of evolutionary dynamics of SP loci across multiple scales. By examining the largest-to-date collection of SP loci in the Enterobacteriales, we showed that our two query locus regions are widespread across the order: *cps* (*kps*) locus region was detected in 22 (11) genera belonging to 5 (3) bacterial families. Importantly, we show that at least one third of all SP locus gene families (LGFs) were detected in multiple species, and that most common gene families showed evidence of frequent jumps between species and locus families. This promiscuous distribution of SP LGFs is consistent with our observation of many cases of horizontal transfer of whole SP loci between bacterial lineages, species and genera. In addition, we found frequent alterations of the LGF content of the SP loci within closely related bacterial lineages, often through gain/loss of a small number of genes. Altogether, our study on the global SP diversity in Enterobacteriales provides robust evidence for SP loci being a major evolutionary hotspot, complementing evidence from previous separate reports on various members of this and other bacterial groups ^51, 55, 57, 58^.

The inter-species view of SP genetics also highlights the range of possible factors that can shape the diversity and distribution of bacterial SPs. It is by now widely accepted that the SP loci are, at least in some bacterial populations, under strong diversifying selection ^33^. Indeed, diversifying selection is consistent with the enormous observed and predicted reservoirs of genetic diversity at SP loci (Fig. S3) and high rates of HGT as it would favour novel locus types arising over time, for example via negative frequency-dependent selection. However, such form of selection would not explain the presence of identical or near-identical loci in distant genomes where we found evidence of no recent locus exchange (Fig. 3C). Such cases, particularly those occurring between divergent taxonomic backgrounds, are suggestive of a form of stabilising selection that preserves the genetic structure of the SP locus over long periods of time. This could occur via positive frequency-dependent selection when some locus types are strongly favoured in certain ecological niches. A notable example are O-antigens in *S. enterica* ^59^ that exhibit high antigenic specificity for particular hosts or environments as they provide protection from predatory protists; incidentally, *S. enterica* was a species in which we detected one of the strongest signals of *cps* locus preservation (c.f., Supplementary Figure S17). However, there might be also other reasons for finding near-identical but diverse locus structures in diverse genomes, including mechanistic factors (e.g., ecological separation or physical barriers to recombination) or simply chance, and the role of such factors cannot be excluded, particularly over the shorter time scales.

Notably, our pan-order study showed that evolutionary dynamics shaping SP loci varied substantially between different bacterial populations, which may be related to the different ecologies exhibited by different members of the Enterobacteriales, thus expanding the scope of previous observations within a single species ^60^. The datasets currently available are not sufficient in size or ecological data/sampling to delve into the specific relationships between ecology and SP locus dynamics in Enterobacteriales. Specifically, we observed a large degree of environmental heterogeneity across the order phylogeny (Supplementary Figure S23) as well as heterogeneity within clonal groups (Supplementary Figure S24), suggesting a considerable degree of transmission between different environmental classes. This implies a substantial sampling bias towards human strains (45% of all isolates were classified as of human origin; see Methods), which in turn makes it difficult to address any ecology-related hypothesis using the studied dataset. Nevertheless, our study not only highlights the importance of ecology in SP locus evolution, but also provides an initial quantification of the effect of evolutionary processes like gene exchange and retention in shaping bacterial gene repertoires, and sets out an approach that could be applied in future to studying more directly the interaction between bacterial ecology and evolution.

Our approach has several important caveats, which we deemed to address in our methodology. First, while we imposed multiple measures to control for the quality of the analysed genome assemblies (genome-based taxonomic assignment, discarding of low-quality genome assemblies and those with low quality SP locus assemblies), it is still conceivable that the dataset includes assemblies that result from mixed or contaminated genome data. Hence we have taken a ‘safety in numbers’ approach, whereby all conclusions drawn from the obtained data are made based on statistical trends, not individual observations. Second, the genome set that we sourced from GenBank includes a mixture of single isolates and collections, sequenced for a variety of reasons and using various different sampling strategies. To mitigate the effect of sampling bias on our estimated parameters, we attempted to minimise it by down-sampling highly similar groups of isolates to single representatives, and estimated parameter uncertainty through resampling isolates with replacement (see Methods). Finally, using *cps* and *kps* loci as models for genetic architecture of SP loci has a disadvantage of missing potentially novel regions of SP diversity. It is possible, even plausible, that some Enterobacteriales species carry other SP-encoding biosynthetic gene clusters that have not been well characterised (for example Villa and colleagues reported a novel putative capsule cluster in a *K. pneumoniae* strain^61^). Hence, this study should not be considered as an exhaustive screen of SP genetic diversity across Enterobacteriales, but rather the largest to date analysis of the diversity and population structure of two relatively well-studied SP locus regions.

Altogether, our study paints a picture of the evolutionary process shaping SP diversity in Enterobacteriales. It provides support for the idea that SP biosynthesis loci are diversity-generating machines, optimised to produce novel phenotypes while minimising the fitness cost of producing sub-optimal combinations ^33^. Diversity generated in such recombination hotspots is subject to action of various evolutionary forces that may differ from those shaping the structure of the underlying bacterial populations within which they occur. Encapsulation has been previously associated with an increased environmental breadth^62^ and increased rates of genetic exchanges ^63^. Our data provide clear evidence that SP loci are also hotspots for HGT at multiple levels, including transfer of whole loci and of subsets of component genes, between lineages of the same species but also across species and genus boundaries, consistent with the action of elevated diversifying selection when compared to the rest of the genome. However, we also show that this is balanced by strong evolutionary constraints on SP loci, which we detect in the form of co-evolution of individual component genes, as well as long-term preservation of similar locus structures in distantly related backgrounds. SPs are at the forefront of most bacterial encounters with novel environmental challenges (e.g., host immunity, microbial or bacteriophage communities). It can thus be expected that keeping up with these challenges requires an unusually high genetic flexibility, as exhibited by SPs, to be able to rapidly generate novel types.

## Methods

### Isolate collection and species assignment

The main dataset consisted of 27,476 genomes from NCBI RefSeq (April 2018) belonging to the 53 genera defined as Enterobacteriaceae by the UK Standards for Microbiology Investigations ^64^: *Arsenophonus*, *Biostraticola*, *Brenneria*, *Buchnera*, *Budvicia*, *Buttiauxella*, *Calymmatobacterium*, *Cedecea*, *Citrobacter*, *Cosenzaea*, *Cronobacter*, *Dickeya*, *Edwardsiella*, *Enterobacter*, *Erwinia*, *Escherichia*, *Ewingella*, *Gibbsiella*, *Hafnia*, *Klebsiella*, *Kluyvera*, *Leclercia*, *Leminorella*, *Levinea*, *Lonsdalea*, *Mangrovibacter*, *Moellerella*, *Morganella*, *Obesumbacterium*, *Pantoea*, *Pectobacterium*, *Phaseolibacter*, *Photorhabdus*, *Plesiomonas*, *Pragia*, *Proteus*, *Providencia*, *Rahnella*, *Raoultella*, *Saccharobacter*, *Salmonella*, *Samsonia*, *Serratia*, *Shigella*, *Shimwellia*, *Sodalis*, *Tatumella*, *Thorsellia*, *Trabulsiella*, *Wigglesworthia*, *Xenorhabdus*, *Yersinia* and *Yokenella*. Note that the definition of Enterobacteriaceae has now been updated to include a subset of these genera, with the rest assigned to new families within the order Enterobacteriales ^38^; hence our analysis can be considered a screen of Enterobacteriales, which uncovered SP loci in the related families Enterobacteriaceae, Erwinaceae and Yersinaceae. False species assignment was corrected using BacSort (github.com/rrwick/Bacsort) – a method that constructs a neighbour-joining tree of all isolates and manually curates monophyletic clades at the species level. Genetic distance was calculated as one minus average nucleotide identity (1-ANI) for all pairs of genomes, where ANI was estimated using kmer-db ^65^ with ‘-f 0.02’ option (which, for a genome size of 5Mb, corresponds to Mash ^66^ with sketch size 10^5^), following by neighbour-joining tree construction using rapidNJ ^67^. We removed (i) isolates belonging to genera of *Arsenophonus* and *Sodalis* as these genera were rare and did not form monophyletic clades (*n* = 6); (ii) isolates with a temporary genus name *Candidatus* that could not be curated using BacSort (*n* = 6); (iii) isolates which could not be assigned to any of the 53 genera. The resulting 27,383 isolates were classified into 45 genera, and assigned into 39 monophyletic genus-groups with the following joint groups: *Buchnera/Wigglesworthia*, *Erwinia/Pantoea*, *Escherichia/Shigella*, *Klebsiella/Raoultella*, *Proteus/Cosenzaea* and *Serratia/Gibbsiella*. For some isolates, especially those descending from rare species in the dataset, species name-reconciliation was problematic. Hence, new species categories (i.e., operational taxonomic units) were defined based on the structure of the distance tree. Monophyletic species groups retained the original species names (e.g., *Klebsiella pneumoniae*), while polyphyletic groups within a genus were split into monophyletic clades with a new unique name (e.g., *Citrobacter unknown* C1; in this paper we refer to these as species-groups, genome assignments are given in Supplementary Table S1). The remaining isolates (*n* = 52) were assigned to the category ‘Other’.

### Quality control of assemblies

Genome assemblies submitted to RefSeq undergo quality control to avoid inclusion of poor quality sequence data in the NCBI public repository ^68^. However, to minimise the risk of finding artificial SP loci due to potential misassemblies, we carried out a further quality check based on consistency of genome summary statistics within groups of related isolates. First, we used hierarchical clustering with complete linkage based on whole-genome distances and a distance cut-off of 0.15 (corresponding to an approximate inter-genus difference) to group all isolates into clusters. Then, within all clusters with 20 isolates or more, we investigated the total genome length distribution and the GC-content distribution, looking for any isolates found outside the 99% confidence interval of these variables assuming they are normally distributed. Any outliers (i.e., genomes that were unusually long/short and that had unusually high/low GC content) were then removed provided that the assembly quality metrics fulfilled the following conditions: *N*50 *<* 10000 and *N*_contigs_ *>* 800. All isolates belonging to the remaining, small clusters were pooled together and their GC-content and total genome length plot revealed two subgroups of isolates (see Supplementary Figure S22). The first group comprised only high quality (completed) genomes from the *Buchnera/Wigglesworthia* genus-group, the small size of which is expected due to ancestral genome reduction in these host-associated organisms ^69^. The second group was screened in the same way as individual clusters before but instead looking at outliers from the non-parametric 95% confidence interval. However, none of these outliers exhibited poor quality assemblies assuming the definition above, and all were kept. In addition, we also tested for potential contamination in the genomes (which could potentially lead to false-positive recombination events) by using CheckM ^70^. We used the same criteria for genome inclusion as those defined for the Genome Taxonomy Database ^71^, namely the completeness estimate *>* 50%, the contamination estimate *<* 10%, and the quality score, defined as completeness *-* 5 *×* contamination, *>* 50. By default, we searched for a CheckM run output at gtdb.ecogenomics.org and, in the case of its absence, we ran CheckM with the taxonomy wf order Enterobacteriales option. Altogether, 49 genomes did not fulfil the criteria above, hence producing a dataset of 27,334 isolates.

### Environmental assignment

Many NCBI RefSeq isolates come from the NCBI Pathogen Detection database, which stores environmental metadata of the bacterial isolates submitted by the users. These include “isolation type” (clinical or environmental/other), “host” (host species) and “isolation source” (e.g., water, faeces, bladder). We used these parameters to classify isolates into three environmental classes: human, animal or non-animal, otherwise classifying them as “unknown” (information missing) or “other” (unfulfilled classification criteria). To classify an isolate into the human class, the “host” category had to match *homo sapiens* or *human*. To classify an isolate into the animal class, the “host” category had to belong to one of the 322 species names listed in a separate file host-names-animal.txt (available via figshare.com; see below). To classify an isolate into the non-animal class, the “isolation source” category had to contain an “env” string. Using these criteria, we found 12,383 isolates of human origin, 4,269 isolates of animal origin, 499 isolates of non-animal origin; 8,551 did not fulfil the classification criteria and 1,632 isolates were missing environmental data altogether. The list of all isolated with the corresponding environmental classes can be found in the Supplementary Table S3.

### Detection of SP loci

#### The *cps* locus

The *cps* locus was defined by the presence of flanking genes *galF* and *gnd* and/or *ugd*, and is associated with the synthesis of group 1 and 4 capsules ^48^ (see example in Figure 1A). While this does not span the full *cps* locus in all species (e.g., in *Escherichia coli* the *cps* locus continues until the housekeeping histidine complex ^48^), the *galF/gnd/udg* region constitutes a major part of the genetic diversity in the *cps* locus region in Enterobacteriaceae. A *cps* locus reference database was created by combining reference loci previously identified in *Escherichia* ^45, 72^ and *Shigella* ^43^ (typically called O antigen loci), *Salmonella* ^44^ (O antigen loci), and *Klebsiella* ^46^ (K antigen loci), amounting to 348 reference locus sequences (duplicate loci containing identical sets of homologous genes were removed). Next, all 27,334 isolates were scanned for the presence of a *cps* locus using an approach similar to the one previously described ^46^. Briefly, for each assembly, we searched for a best match among all reference sequences using blastn. Given such best match, we defined *C* (percentage of the best match found in the assembly), *Nm* (number of best match genes missing in the detected locus), *Ne* (number of extra genes present in the detected locus), and *Nc* (number of contigs on which locus was found). The best match was defined as a good match if any of the following conditions was met:

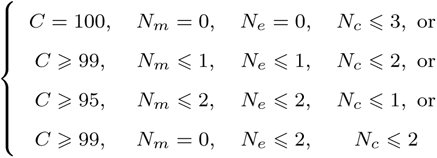

If the condition was not met but *Nc* = 1, we extracted the locus and reannotated it using Prokka ^73^ with a custom gene database. The locus was defined as missing if (i) the only genes detected were core genes (*galF/gnd/ugd*) or other sugar-synthesis genes (*rmlACBD, manBC, fcl, gmd, galE, glm, rmd*); (ii) if there was a single contig with only core genes; (iii) if no core genes were found. In the first round, all new loci present in a single contig were extracted, and loci without gene duplications or stop codons and with all core genes present were added to the reference database, altogether comprising 994 reference sequences. In the second round, all loci which had a good best match, as defined above, or new loci in a single contig (*Nc* = 1) were extracted. Loci had transposons removed using ISFinder (e-value threshold of 10*^-^*^20^), and frameshift mutations corrected using Decipher R package^74^. According to those definitions, 3,275 isolates were missing the *cps* locus, and 18,401 had the locus, which was then extracted.

#### The *kps* locus

The *kps* locus was defined by the presence of flanking genes *kpsF, kpsE, kpsD, kpsU, kpsC, kpsS, kpsT* and *kpsM* (group 2 capsules) and *kpsD, kpsM, kpsT, kpsE, kpsC* and *kpsS* (group 3 capsules), as reviewed in ^48^ (see example in Figure 1A). The initial reference sequence database consisted of six sequences from *Escherichia coli* ^75^. Locus extraction was performed in the same way as with the *cps* locus, except that the locus was defined missing in the case of absence of any of the three genes *kpsM, kpsT, kpsC*, and the extended reference database had 106 sequences. With those definitions, 24,759 isolates were missing the *kps* locus, and 2,356 had the locus extracted.

### Locus gene families and locus types

Coding sequences within all detected loci were clustered into locus gene families (LGFs), separately for the *cps* and *kps* locus regions. The clustering was done using mmseqs2 ^76^ with default settings and the 50% sequence identity threshold at the protein level (‘--min-seq-id 0.5’ option). For each pair of loci from isolates *i*1 and *i*2, a Jaccard distance (*J*) was calculated between the SP loci in both isolates as

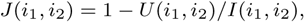

where *U* (*i*1*, i*2) is the number of LGFs in a union of all LGFs in both loci and *I*(*i*1*, i*2) is the number of LGFs in an intersect of all LGFs in both loci. Locus types were defined by connected components of a locus similarity network clustered at *J* 6 0.1. A representative set of loci for all Enterobacteriales was chosen as single representatives of all loci within each species. This led to a set of 2,654 *cps* loci and 332 *kps* loci (the set used in Fig. 2).

### Definition of within-species lineages

We defined bacterial lineages *Lx* by using a hierarchical clustering approach with complete linkage at the whole-genome distance threshold *x*. Depending on the analysis, we used two different lineage definition thresholds, *x* = 0.004 and *x* = 0.01 (note that an evolutionary distance cut-off of 5% has been proposed as a criterion for delineating bacterial species ^77^).

### Detection of ribosomal sequence types (rST)

We identified ribosomal sequence types (rST) for each isolate in reference to a standard typing scheme ^78^, using a custom procedure implemented in a R script. First, we used blastn (with megablast option) to search all unique ribosomal sequences similar to the reference alleles of the 53 ribosomal gene loci with an e-value threshold of 10*^-^*^10^ in each genome assembly, with the top hit for each locus recorded. Each isolate was then assigned a combination of 53 numbers, corresponding to an allele index at each ribosomal gene. Isolates with more than five missing alleles were regarded as with an unknown rST. Finally, isolates with identical allele combinations (disregarding the missing loci) were clustered into rSTs. The detected rSTs were used to define representative datasets by: (i) identifying unique combinations of rST and locus type *Ci* (for *i* = 0, 1 or 2), and (ii) for each unique rST/locus type combination choosing the isolate with the greatest *n*_50_ value.

### Identification of recent locus exchanges in pairs of genomes

We aimed to detect recent locus exchanges between pairs of genomes sharing locus types (using the *C*1 definition). A simple pairwise genome search yielded over six million pairs sharing the locus type in the *cps* region. As the large majority of these pairs were isolates belonging to the same clonal family (and thus most likely sharing the locus through common ancestry), we filtered this dataset in two steps. First, we only considered genome pairs belonging to different lineages (under the *L*0.004 definition). Second, we only considered a single representative of a combination of rST and a *C*1 locus type (as defined above). This resulted in 7,000 isolates forming 99,177 pairs, and in 1,114 isolates forming 23,928 pairs for the *cps* and *kps* loci, respectively. For each pair, we then downloaded predicted coding regions in the relevant assembly from NCBI RefSeq ^79^ and, when unavailable, we predicted them using Prodigal ^80^ with default settings, and removed all protein sequences previously found in the *cps* or *kps* regions. Next, we used mmseqs2 with 10*^-^*^50^ e-value threshold to: (a) align SP locus protein sequences in one genome against those in the other genome, (b) all other protein sequences in one genome against those in the other genome. If a protein in one genome gave a hit to multiple proteins in the other genome, only the most similar sequence was retained. Then, we created a distribution of the genome protein sequence comparison by (a) randomly sampling *np* protein sequences (where *np* is the number of homology groups common to both loci), (b) calculating the mean percentage similarity between them, and (c) repeating this process 10,000 times. The locus type was considered recently exchanged via recombination if the locus proteins were amongst the top 5% of the most similar sets of proteins (see also Supplementary Figure S7). Conversely, if the locus proteins were in the bottom 60% of the most similar sets of proteins or the locus evolutionary distance was greater or equal to the whole-genome locus distance, the locus was considered selectively maintained.

### Quantification of purifying selection

Among all representative isolates (see above), 7,403 isolates had a *cps* or *kps* locus. The coding sequences from those isolates were then clustered using mmseqs with default settings and ‘--min-seq-id 0.5’ option to find family-wide homology groups (HGs). This resulted in a total of 36,019,427 proteins clustered into 28,441 HGs. Gene sequences assigned to each HG were then aligned at the protein level with mafft ^81^ with default settings and reverse aligned with pal2nal ^82^. Maximum likelihood tree for each HG was created using FastTree ^83^ with ‘--gtr --gamma’ options, and midpoint rooted. Ancestral SNPs were recovered using ClonalFrameML ^84^ with the ‘-imputation only’ option, allowing us to recover all independent synonymous and non-synonymous substitutions that occurred over each branch of each HG phylogeny. Next, all proteins were aligned against the representative *cps* and *kps* proteins with mmseqs align with an e-value threshold of 10*^-^*^32^. HGs in which 90% of proteins had hits to one of the protein categories in the reference genome (‘SP locus’ or ‘other’) were assigned to that category, otherwise ignored. Considering only clusters with 100 sequences or more as informative, we obtained 26,231 assigned to the ‘other’ category and 261 HGs were assigned to the ‘SP locus’ category; 73 HGs were disregarded.

### Calculation of jump-rates between bacterial species

All nucleotide sequences assigned to a given locus gene family (LGF) were aligned with mafft with default settings. Next, a maximum likelihood tree was build using FastTree with ‘--gtr --gamma’ options and midpoint rooted. We then used the asr max parsimony() function from R package castor to infer bacterial species as ancestral states at internal nodes of the tree ^85^, and considered resolved states as those with marginal likelihood assigned to a single species of at least 0.95. Next, we considered all tree edges with resolved states at the upstream and downstream nodes, and these were divided into those where there was a change in species along this branch (species jump) and those where there was no change (no jump). The species jump rate was calculated as the ratio of the number of branches with detected species jump to the summed branch lengths of all considered tree edges. To account for uncertainty in tree inference, for each alignment we generated 100 bootstrap trees. To account for uncertainty in the number of sequences within each LGF, for each tree we randomly subsampled (with replacement) branches of that tree 100 times. The species-jump rate was estimated for each LGF tree in the overall 10^4^ bootstrap sample and the median and 95% confidence intervals of the obtained distribution were reported. The locus-wide species-jump rates were calculated in the same way based on concatenated alignments of all LGF present in all isolates for a given SP locus. In that calculation, isolates with less than three low-to-middle frequency genes (20% cutoff) were discarded as in these cases locus-family assignment may have been problematic leading to false-positive locus-family jumps.

### Estimation of SP gene flow within lineages

To quantify SP gene flow within bacterial populations, we first identified, for both *cps* and *kps* locus, a subset of isolates where the locus was identified in a single contig. This was done to avoid mistaking gene absence due to inability to assemble the full locus with gene absence due to actual gene gain or loss. We then considered a dataset of single representatives of the rST and *C*0 locus types. Then, for each lineage defined at the *L*0.004 level, we generated a neighbour-joining tree based on pairwise whole-genome distances using rapidNJ ^67^ as well as a LGF presence/absence matrix. We then considered lineages which contained at least 10 isolates and at least 3 accessory LGFs (i.e., at least three gene families were gained or lost in that lineage). The tree and the LGF presence/absence matrices were then used as inputs for GLOOME ^85^ to infer the gene gains and losses at each branch of the tree using the maximum parsimony model. Co-mobility coefficient of a pair of LGFs, *i* and *j*, was calculated as a co-occurrence of gain or loss events between *i* and *j*, namely as a proportion of a sum of all branch lengths in which *i* and *j* were found to be co-gained or co-lost to the sum of all branches in which *i* or *j* were found to be gained or lost.

### Identification of co-evolving capsule genes

Co-evolution coefficient *CVij* between LGFs *i* and *j* was estimated as 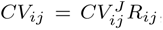, where 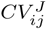 is a co-occurrence coefficient calculated as a Jaccard index of the number of isolates with LGF *i* present and those with LGF *j* present, and *Rij* is a correlation coefficient reported by the Mantel test using matrices of pairwise distances between sequences from co-occurring LGFs. Mantel test calculation was performed using ecodist R package ^86^ using Kimura 2-parameter distance measures.

### Transparency of the study

The data generated in this study (extracted nucleotide and protein locus coding sequence, their assignment into LGFs, genome phylogeny of all isolates and list of NCBI animal hosts) can be found under the following links:

- https://doi.org/10.6084/m9.figshare.8166845
- https://doi.org/10.6084/m9.figshare.8166794
- https://doi.org/10.6084/m9.figshare.8166815
- https://doi.org/10.6084/m9.figshare.11279000
- https://doi.org/10.6084/m9.figshare.11322116

The software used to identify SP locus sequences in *cps* and *kps* regions in Enterobacteriales genomes, as well as all locus reference sequences used in this study, are available at: https://github.com/rmostowy/fastKaptive.

## Glossary

SP: surface polysaccharide
HGT: horizontal gene transfer
LPS: lipopolysaccharide
ANI: average nucleotide identity
CDS: coding sequence
LGF: locus gene family

## Supporting information

Supplementary Tables (ZIP)

## Acknowledgements

This work was supported by a Senior Medical Research Fellowship from the Viertel Foundation of Australia (KEH), the Bill and Melinda Gates Foundation (KEH), the UK Medical Research Council (FL, grant MR/N010760/1), the Imperial College Research Fellowship (RJM) and the Polish National Agency of Academic Exchange (RJM). The authors also acknowledge (i) the support of computational resources of PLGrid Infrastructure, (ii) MRC CLIMB for the use of high-performance computational facilities, (iii) Mat Beale for kindly sharing his scripts on CheckM genome analysis.

