## Supplementary Material (PDF) for "Diversity and evolution of surface polysaccharide synthesis loci in Enterobacteriales"

### Supplementary Text

#### S1. What predicts SP locus diversity

SP locus similarity networks in Figure 1 also clearly show that we found many more locus types in some genera than others. Furthermore, those genera in which most locus types have been found (like *Escherichia*, *Klebsiella* and *Salmonella*) were incidentally the largest isolate collections in our dataset. We thus investigated whether such locus diversity, measured as the number locus types (defined at the $C_1$ level; see Methods), is a consequence of the greater genetic diversity detected in some genera, or simply the greater number of sampled sequences. To this end, we examined which variable best predicts the number of SP loci identified in a defined taxonomic group (genus or species): the number of genomes sampled, or overall nucleotide diversity (evolutionary distance) captured within that sample.

Results are shown in Figure S3A. We found that, for both genus and species level taxonomic groupings, sample size was a much better predictor of antigenic diversity (measured by number of unique locus types in the *cps* or *kps* region) within the group than was genetic diversity captured by the sample (measured by the average root-to-tip distance measured on the ANI distance-based neighbour-joining tree of strains). One consequence of performing this test using common species and genus definitions is that it is based on historically determined taxonomic group definitions, some of which may be much more genetically diverse than others. To overcome this potential bias, we performed a systematic clustering at a given (whole-genome) ANI threshold distance using hierarchical clustering with complete linkage. We then re-examined the same relationships at different thresholds, spanning a wide evolutionary range from genus level (around 85%-90% ANI similarity), to species level (around 95% ANI similarity), to lineage level (around 99% ANI similarity). The results are shown in Figure S3B and indicate that sample size per taxon was a strong predictor of within-taxon SP locus diversity, often explaining at least 50% variance in observed numbers of loci; whereas within-taxon genetic diversity had little effect.

If sample size is a good predictor of locus diversity, it would suggest that many more locus types could be discovered in the future as new isolates are collected and sequenced. We attempted to quantify this by considering an ecological species modelling approach, whereby sampled ribosomal sequence types (rST) were considered ecological "sites", and *cps/kps* locus types were considered observed "species" in those "sites". With this assumption, we calculated the rarefaction curves, i.e., the expected number of "species" (loci) per new rST. This was done using the `iNEXT` package in R[1]. The results for six species with the largest number of locus types are shown in Figure S3C. This model predicts a large reservoir of unobserved antigenic diversity, particularly in *cps*, but that the size of this repertoire could vary between species. This would be consistent with the idea that some enteric species might have a greater population-wide SP diversity than others.

### S2. Comparison of jumping and non-jumping loci

As locus-type sharing between different bacterial lineages was sometimes best explained by recent locus exchange (blue in Figure 5) and sometimes by lack thereof (red in Figure 5), we wanted to examine if a propensity of a locus-type to undergo exchange (to the extent we could detect it) was associated with certain predictors. To this end, we investigated two potentially important factors: frequency and genetic content of the locus.

One hypothesis was that recent exchange was more likely to be detected in more frequent locus-types. Here we did not speculate on the direction of causation (whether HGT was actually driving the changes in locus-type frequencies in the population or simply interacting with positive selection), only that the detected likelihood of locus-type exchange given sharing correlated with the frequency of that locus in the population. To address this hypothesis, we first looked at the relationship between locus-type frequency, measured using the representative collection of isolates mentioned in the main text, and (i) its propensity to undergo recent exchange (proportion of pairs with that locus-type that have recently exchanged that locus) and (ii) its propensity to not undergo recent exchange (proportion of pairs with that locus-type that have not recently exchanged that locus). To minimise the effect of pseudo-replication, we only considered unique combinations of $L_{0.004}$ lineages and a locus type. Results, shown in Table S4, Figure S12A for *cps* locus and Figure S12C for *kps* locus, demonstrate no obvious positive correlation between recent HGT and locus-type frequency, though borderline significant for the *kps* locus. Next, for all isolate pairs, $i$ and $j$, with a locus-type belonging to each of the three classes (jumpers/non-jumpers/unclear), we compared the distribution of $f_i f_j$ between the three classes (where $f_i$ is the frequency of lineage $i$ in a representative dataset), expecting that we would be more likely to detect recent exchanges in more frequent lineage combinations (Figure S12B for *cps* locus and Figure S12D for *kps* locus). Again, in the case of the *cps* locus we rejected the alternative hypothesis that recently exchanged locus-types occurred between more frequent combinations of lineages (Wilcoxon one-sided test, $p = 1$), although we confirmed it for the *kps* locus ($p < 10^{-16}$). These results show that there is no obvious relationship between locus exchange via HGT and locus/isolate population frequency, suggesting that such a relationship is highly non-linear and likely dependent on the multitude of selective and ecological forces interacting with the microbial population.

The second hypothesis we examined was that the propensity to undergo locus exchange by HGT given sharing, or to not undergo such exchange, was driven by specific locus genetic content (namely by carrying genes with a specific function). To address it, for each locus-type we first estimated a probability of exchange or lack thereof given sharing, and asked whether the estimated probability is significantly greater than expected compared to these probability values on a population-wide scales (i.e., across all locus-types). We calculated such significance using a binomial proportion test with a confidence interval $1 - p_{\text{thr}}$, where $p_{\text{thr}} = 0.05/n$ is the Bonferroni corrected p-value threshold accounting for multiple testing amongst $n$ locus-types. As seen using a subset of the most frequent locus types shown in Figure S13A (those shared by at least 100 pairs), some locus-types were indeed highly likely to be exchanged by HGT while others were highly likely to not have been recently exchanged. As expected, we also found that the two measures were strongly negatively correlated (Figure S13B; Kendall's rank correlation, *cps*: $\tau = -0.69$, $p = 3.2 \times 10^{-13}$, *kps*: $\tau = -0.82$, $p = 3.6 \times 10^{-4}$). We then defined a group of extreme jumpers (probability of recent exchange given sharing significantly greater than 0.9 and $p < p_{\text{thr}}$) and a group of extreme non-jumpers (probability of absence of recent exchange given sharing significantly greater than 0.9 and $p < p_{\text{thr}}$), and asked wether there were any locus gene families (LGFs) that were significantly over- or underrepresented in jumpers (non-jumpers) given their presence/absence pattern amongst the jumpers (non-jumpers) vs all other locus-types. We calculated odds ratios for each LGF with confidence intervals for the Bonferroni-corrected significance levels $p_{\text{thr}} = 0.05/n$ where $n$ is the number of all unique LGFs considered in a given class. All genes putatively associated with jumpers/nonjumpers (i.e., those with $p < 0.01$) are shown in Figure S13C, and their functional annotations are shown in Table S5. We found no genes putatively associated with jumpers, and all but one genes putatively associated with non-jumpers were non-significant when correcting for multiple testing. The only significant result was obtained for HG0299, which is a distant copy of the polysaccharide export protein *kpsE*, typical of the group 2 and 3 capsules. We also found three genes with relatively low p-values (though still non-significant) that were putatively associated with non-jumpers, namely HG3110 and HG2781 (*wzm* and *wzt*, both ABC transporters), and HG3949 (*wfdA*, a glycosyltransferase). (See Figure S13 and Table S5.)

### Supplementary Figures

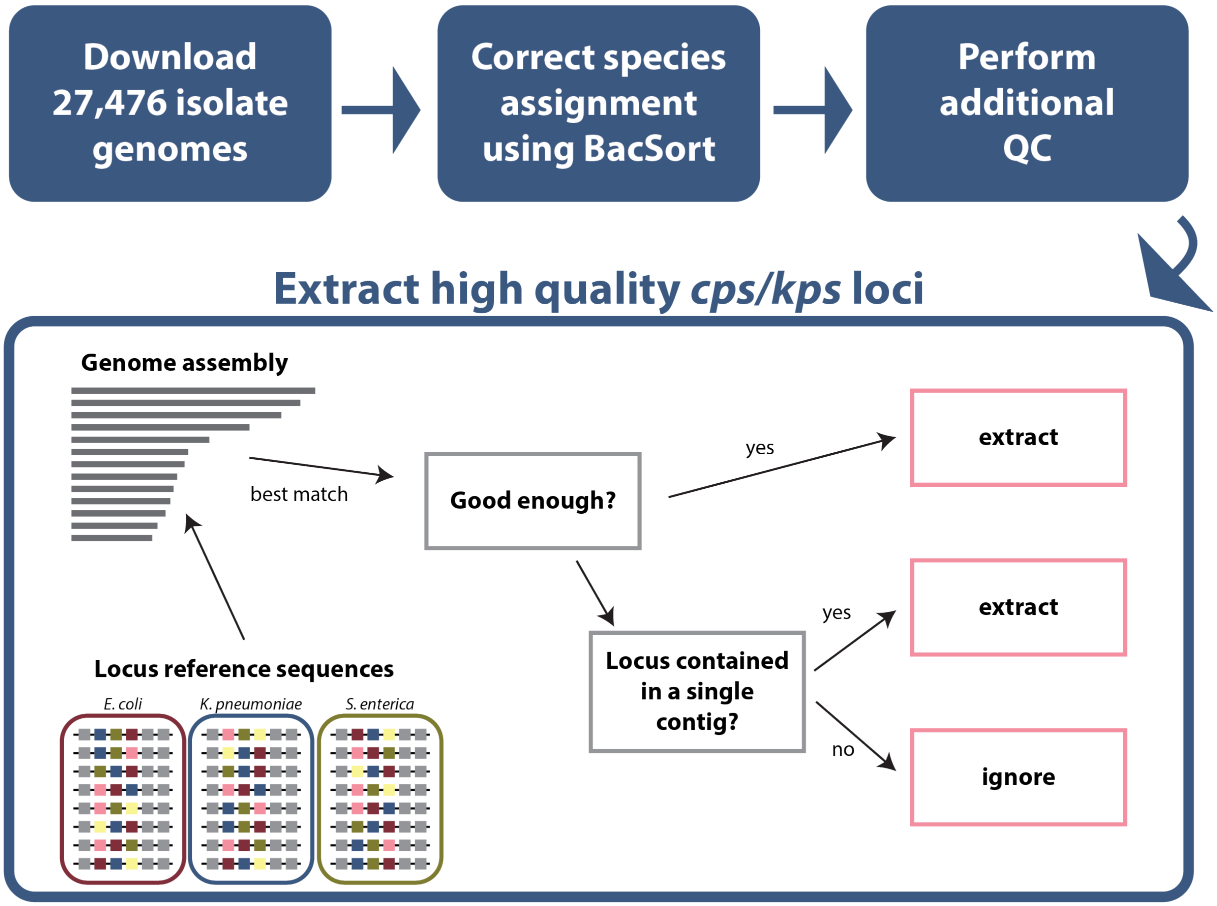


**Figure S1.** Workflow summary for mining 27,476 bacterial isolate genomes for the presence of *cps* and *kps* locus, as described in Methods in detail.

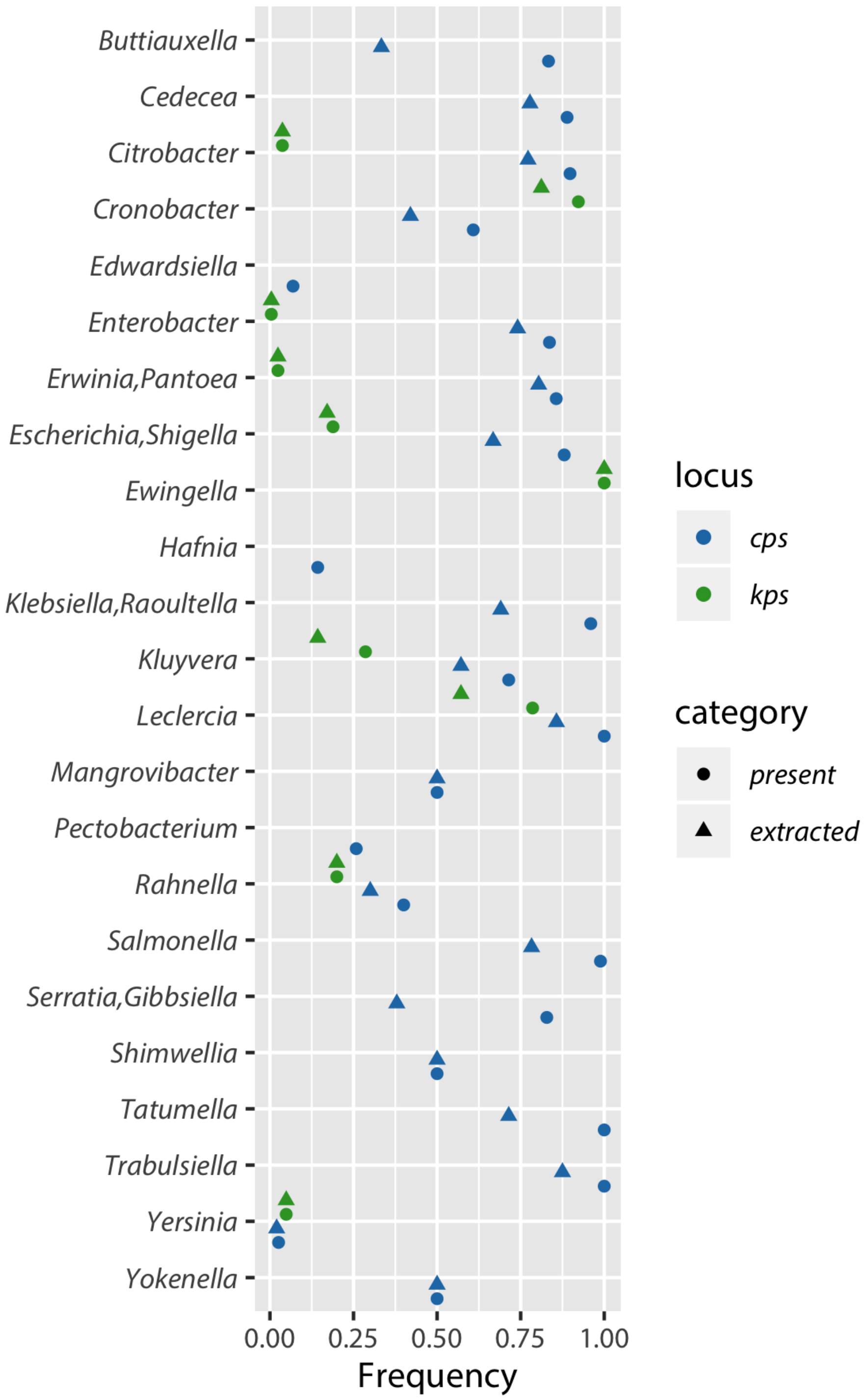


**Figure S2.** Frequency of the locus region *cps* (blue) or *kps* (green) in different genera. The frequency was calculated as a proportion of isolates within each genus-group in which the locus was detected as present (circle) and those in which it was of enough high quality to be extracted (triangle).

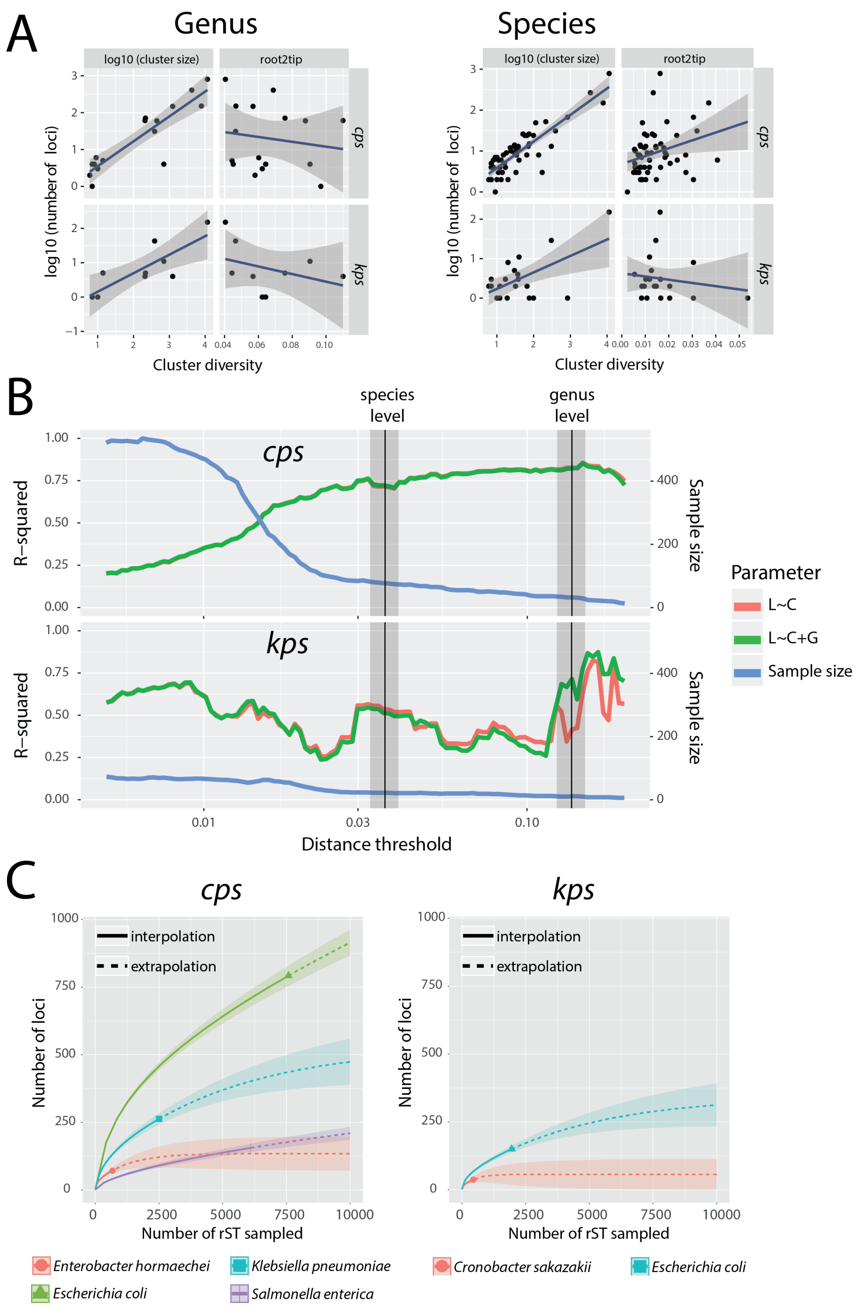


**Figure S3. What predicts SP locus diversity?** (A) Number of SP locus types per genus (left) and species (right) as function of the sample size (log of cluster size) and diversity (measured by the average root-to-tip distance of the neighbour-joining ANI distance tree) for the two locus regions, *cps* and *kps*. (B) All isolates were clustered into groups based on the whole-genome distance threshold, displayed on the X-axis. The blue line shows the sample size, namely the number of these groups (with the minimum size of 5 isolates per group). For a given threshold, we fitted two linear models to explain the locus diversity measured by the number of locus types across different groups ($L$). The first model uses the group size ($C$; log of the number of isolates) as the predictor. The second model uses a combination of the group size ($C$) and genetic diversity ($G$; average root-to-tip distance). The red and the green curves show the $R^2$ obtained from these model fits, respectively. (C) Rarefaction curves for six species of Enterobacteriaceae, predicting the number of locus types per species as a function of the sampled ribosomal sequence types (rSTs).

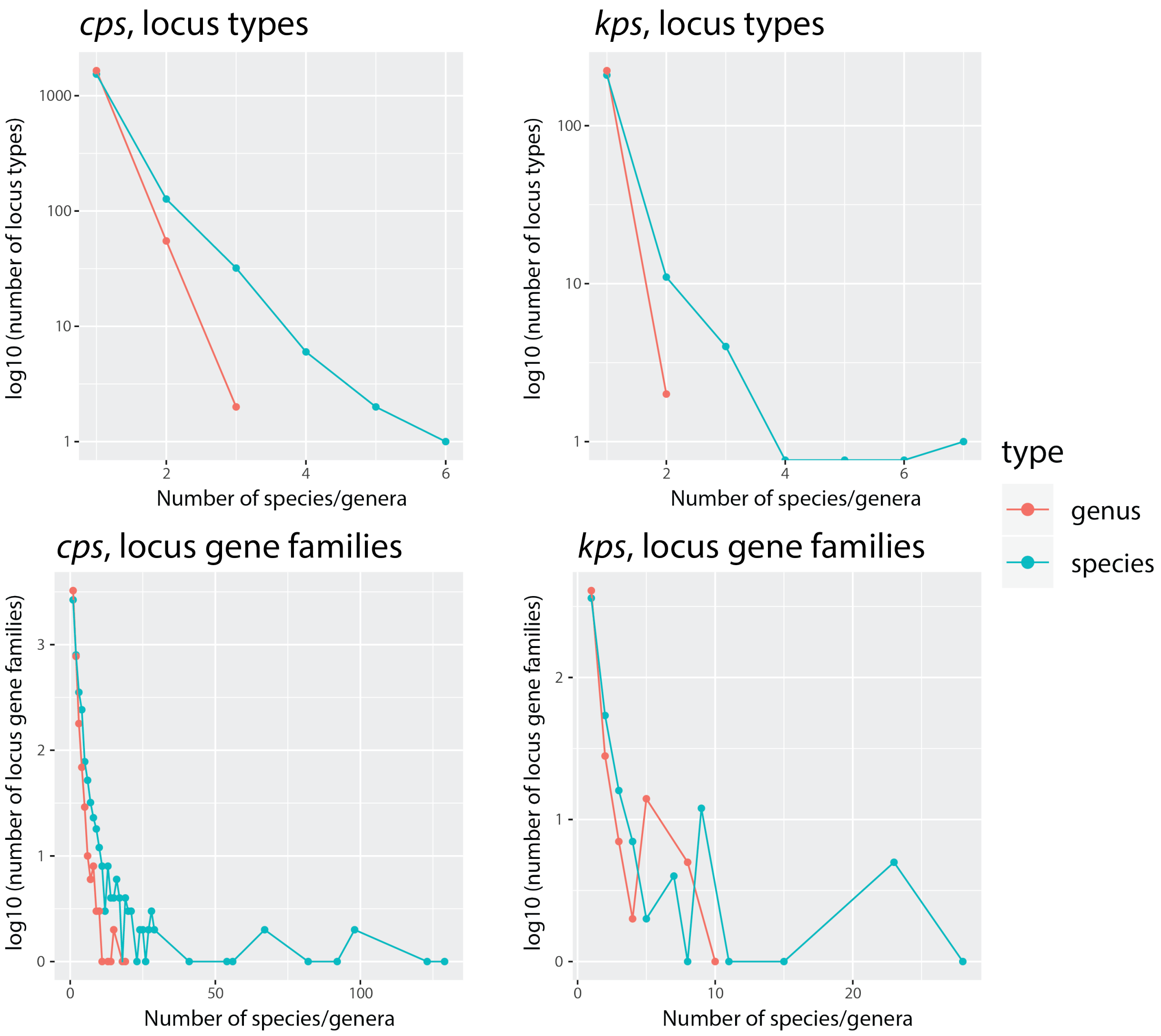


**Figure S4. Distribution of SP loci and genes across species and genera**. The top row shows the frequency distribution of SP locus types across different species (blue) and genera (red) in the *cps* region (left) and the *kps* region (right). The Y-axis shows the log10 value of the number of locus types, and the X-axis shows the number of species of genera. The values on the left-hand side of the X-axis show locus types which are found in a small number of species (majority of locus types), while the large values on the X-axis show locus types which are found in multiple species and genera (minority). The bottom row shows the same distribution for the locus gene families identified in the *cps* region (left) and the *kps* region (right).

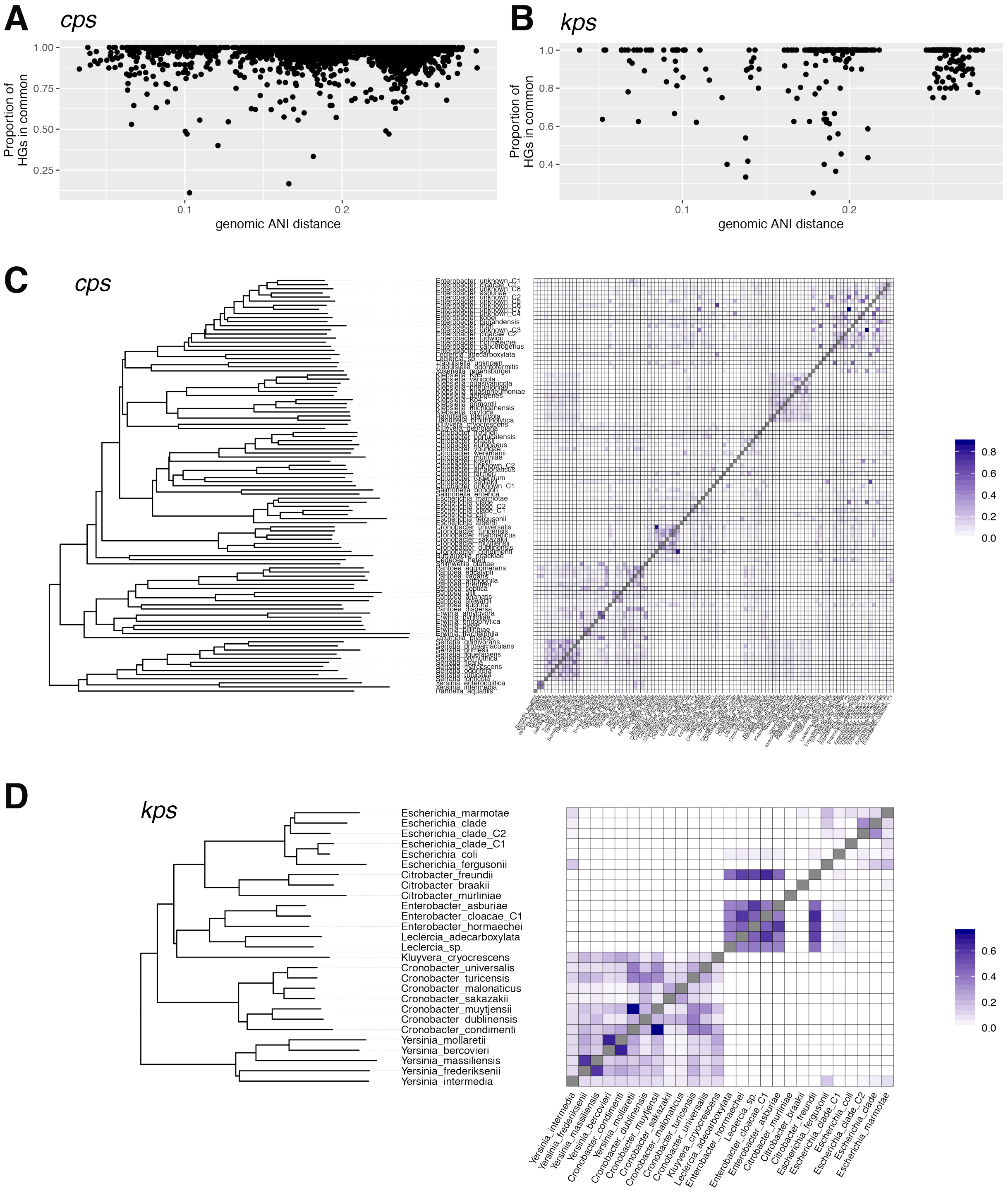


**Figure S5. Species-level relation between locus and genome population structure.** (A) Each point corresponds to a pairwise comparison between genus-groups for the *cps* and *kps* locus. The X-axis represents the genomic ANI distance between representative members of each genus. The Y-axis represents the Jaccard distance between the unique sets of homology groups identified in each genus. All homology groups present in 20% or more of all representative isolates were discarded in this calculation of the Jaccard distance. (B) Same but for the *kps* locus region. (C) SP locus LGF sharing patterns between species, in complete analogy to Figure 2C for the *cps* region, and (D) for the *kps* region.

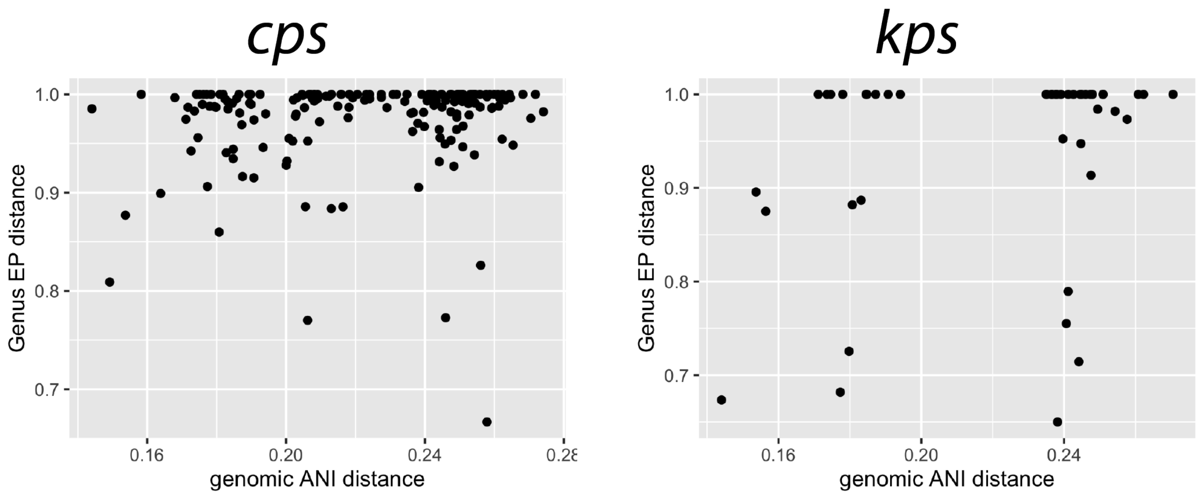


**Figure S6. Weak evidence for correlation between genomic and SP locus population structure at the genus level**. Each point corresponds to a pairwise comparison between genus-groups for the *cps* and *kps* locus, as shown in Figure 2. The values are obtained in the same way as in Figure S5.

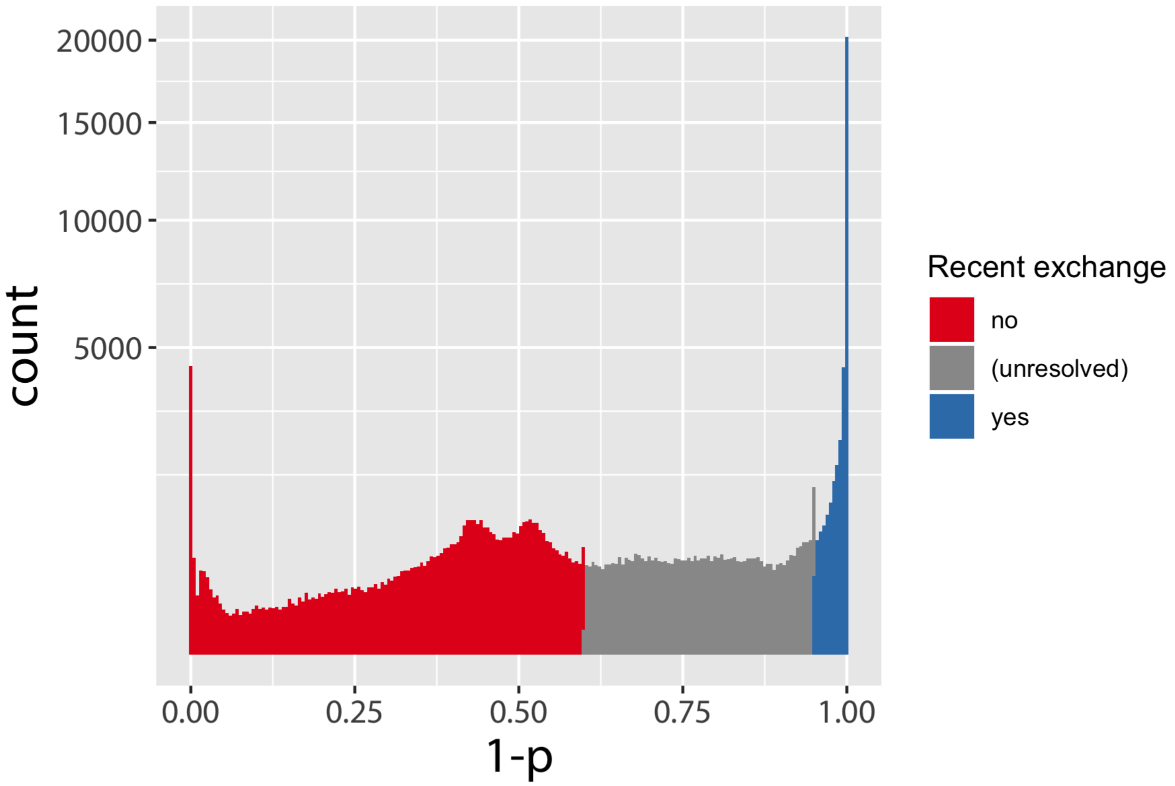


**Figure S7. Distribution of p-values in the recombination test**. Histogram shows the distribution of $y_{ij} = 1 - p_{ij}$, where $p_{ij}$ is the p-value obtained in the test for locus exchange between isolates $i$ and $j$. Results for *cps* and *kps* are combined. Locus sharing via recombinational exchange was called when $p_{ij} < 0.05$, namely in those cases where in 95% of comparisons between isolates $i$ and $j$ the locus was more genetically similar than other coding sequences (blue bars). Cases with $p_{ij} \geqslant 0.4$ were considered to provide evidence for the absence of recombination, hence suggesting sharing via selective maintenance (red bars). For the remaining cases, there was not enough statistical power to claim one or the other cause of sharing (green cases).

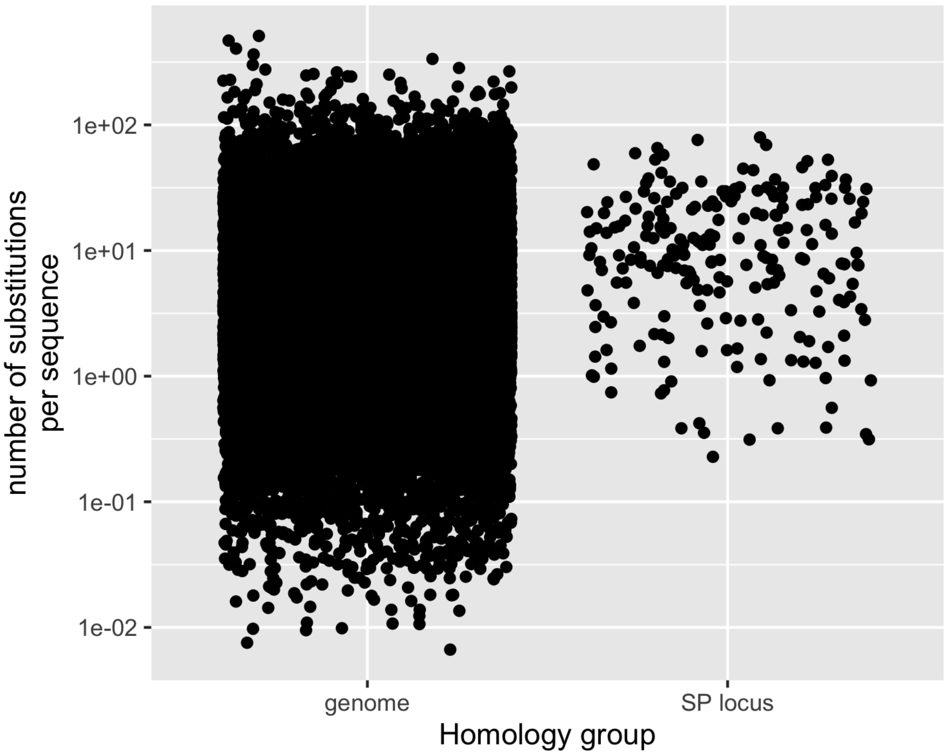


**Figure S8.** Estimated conservedness of HGs associated with genome (left) and SP locus (right), as defined in Figure 3E. Conservedness was calculated as the number of independent substitutions per sequence. We found that SP locus HGs were less conserved that the genome HGs (Student's t-test $p = 1.23 \times 10^{-7}$).

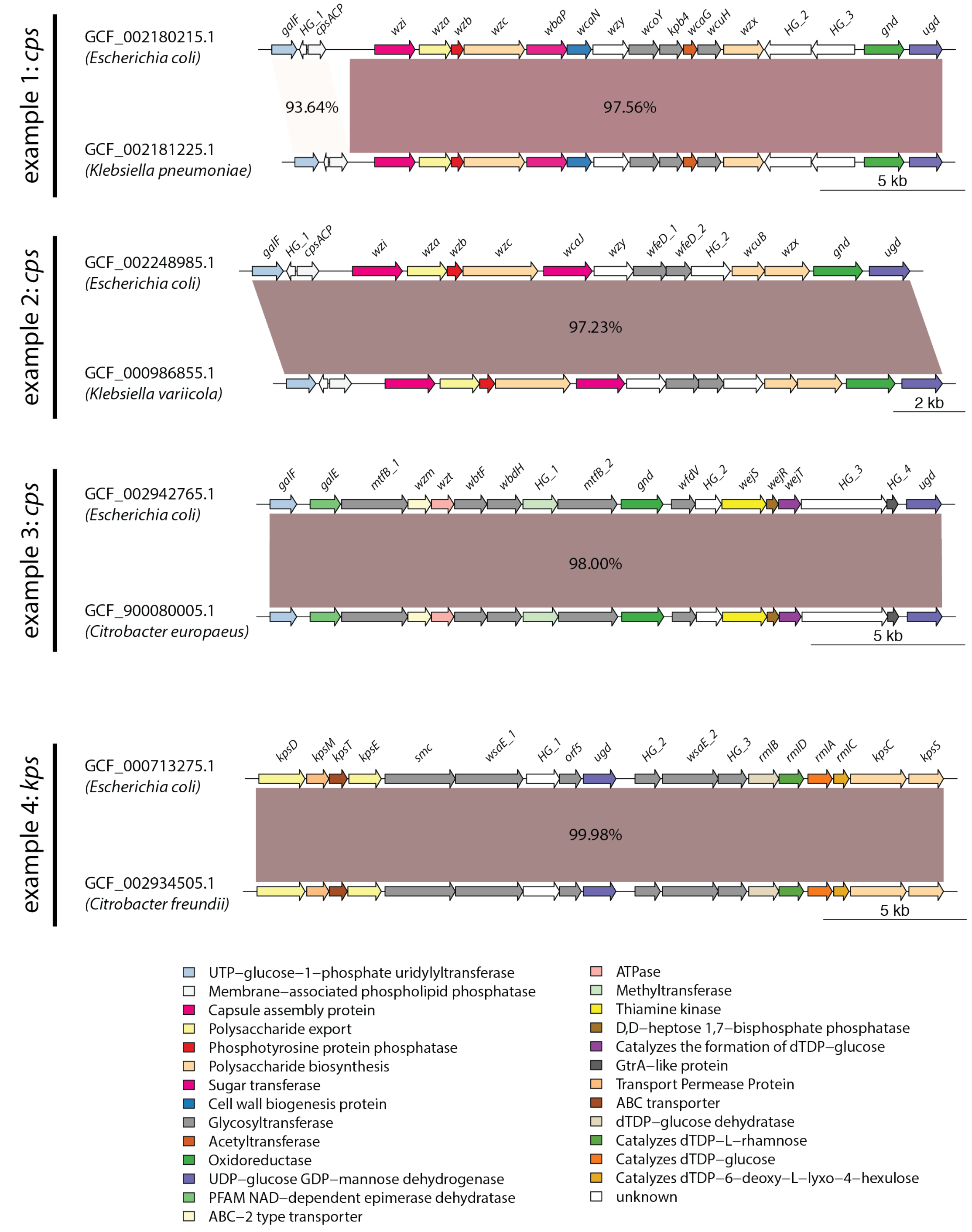


**Figure S9.** Four examples of horizontal locus exchange between species (i.e., where SP locus distances rank in the lowest 5% of gene distances for a pair of lineages, and thus are inferred to have been exchanged long after the lineages themselves diverged), including *cps* (top three) and *kps* (bottom). Each alignment depicts the locus genetic architecture and the percentage identity between aligned regions. Colours correspond to gene functions as predicted by `eggNOG-5.0` (`http://eggnog-mapper.embl.de`).

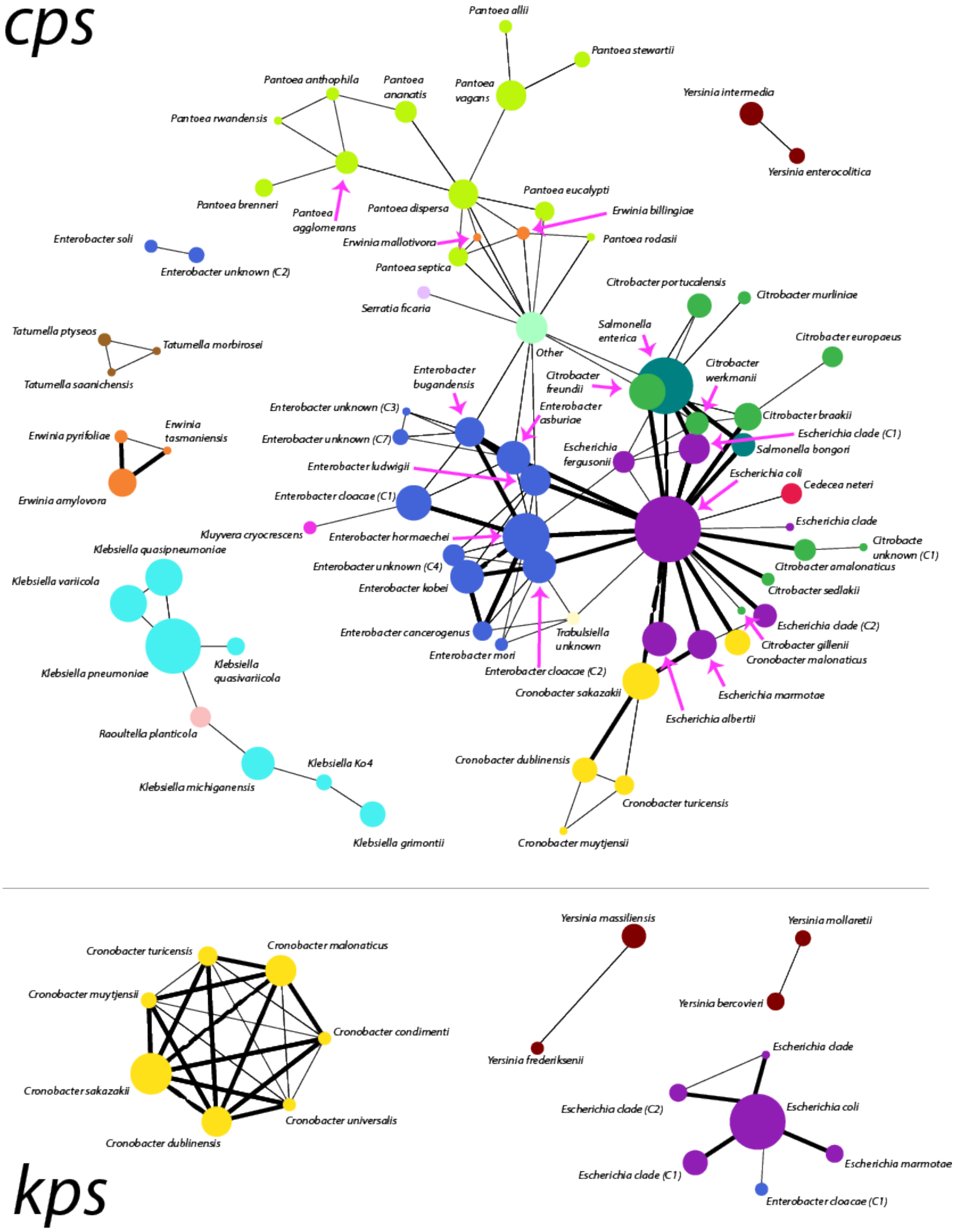


**Figure S10. Network of SP locus type sharing in the absence of recombinational exchange**. The network was constructed in the same way as 4, but with edges linking species-groups for which there was evidence of locus type sharing in the absence o recent exchange (red points in Figure 3).

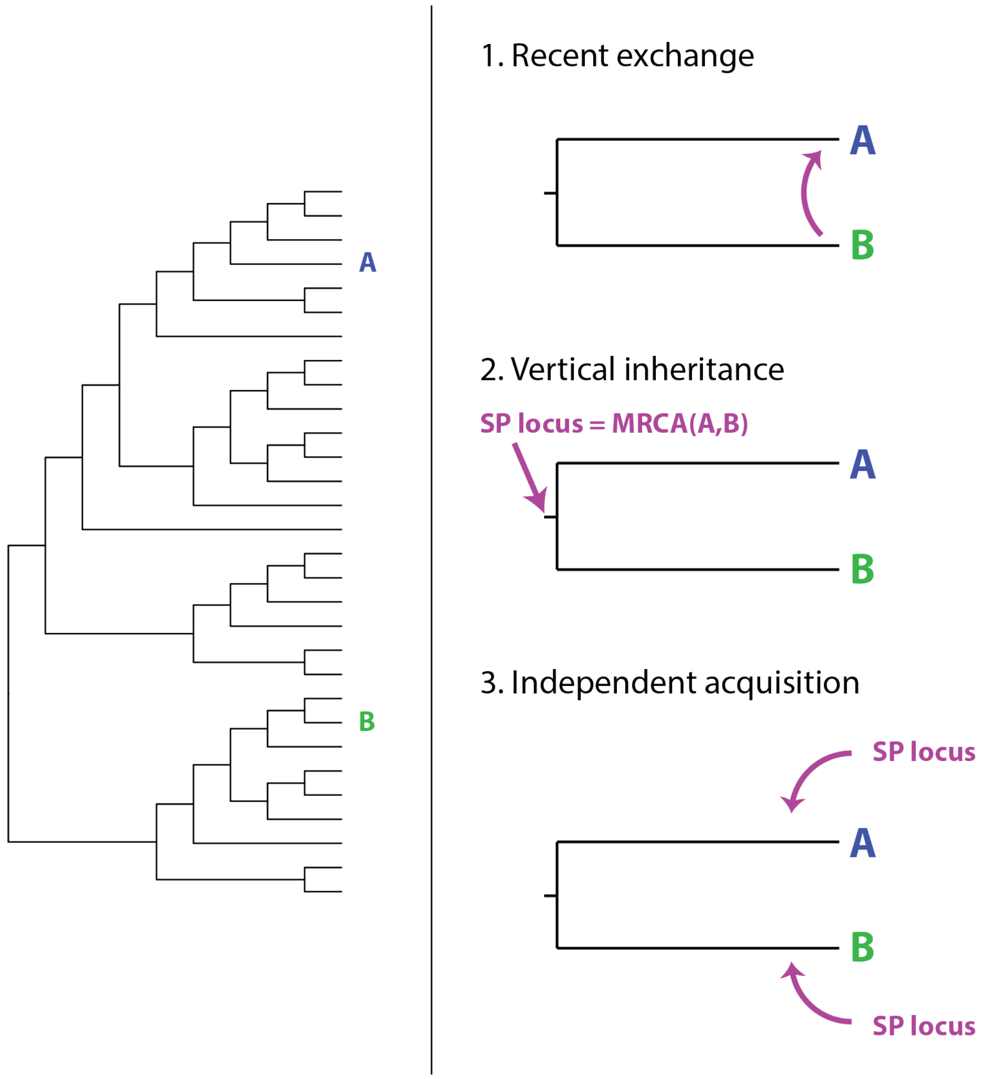


**Figure S11.** Three scenarios to explain SP locus type sharing (i.e., presence of highly similar sets of SP locus genes) in bacterial genomes from different taxonomic backgrounds. The tree on the left represents a structure of a population from which two isolates, A and B, are sampled. These isolates share a locus type, and three potential scenarios that have led to this are shown on the right hand side. Under scenario 1, A and B share a locus type as a result of a recent horizontal exchange of the locus between them (blue categories in Figures 3 and 5). Under scenario 2, A and B share a locus type as a result of vertical inheritance of that locus since the most recent common ancestor of A and B; the most recent common ancestor of SP loci in A and B was carried by the most recent common ancestor of A and B. Under scenario 3, A and B share a locus type as a result of multiple acquisitions of that locus (either via whole locus exchange or gene reassortment) from another source (or multiple sources). Under a neutral model, the likelihood of scenarios 1 and 2 are expected to decrease as a function of genetic distance between A and B, while the likelihood of scenario 3 is expected to remain constant as acquisition of the locus type by lineage A is independent from acquisition by lineage B.

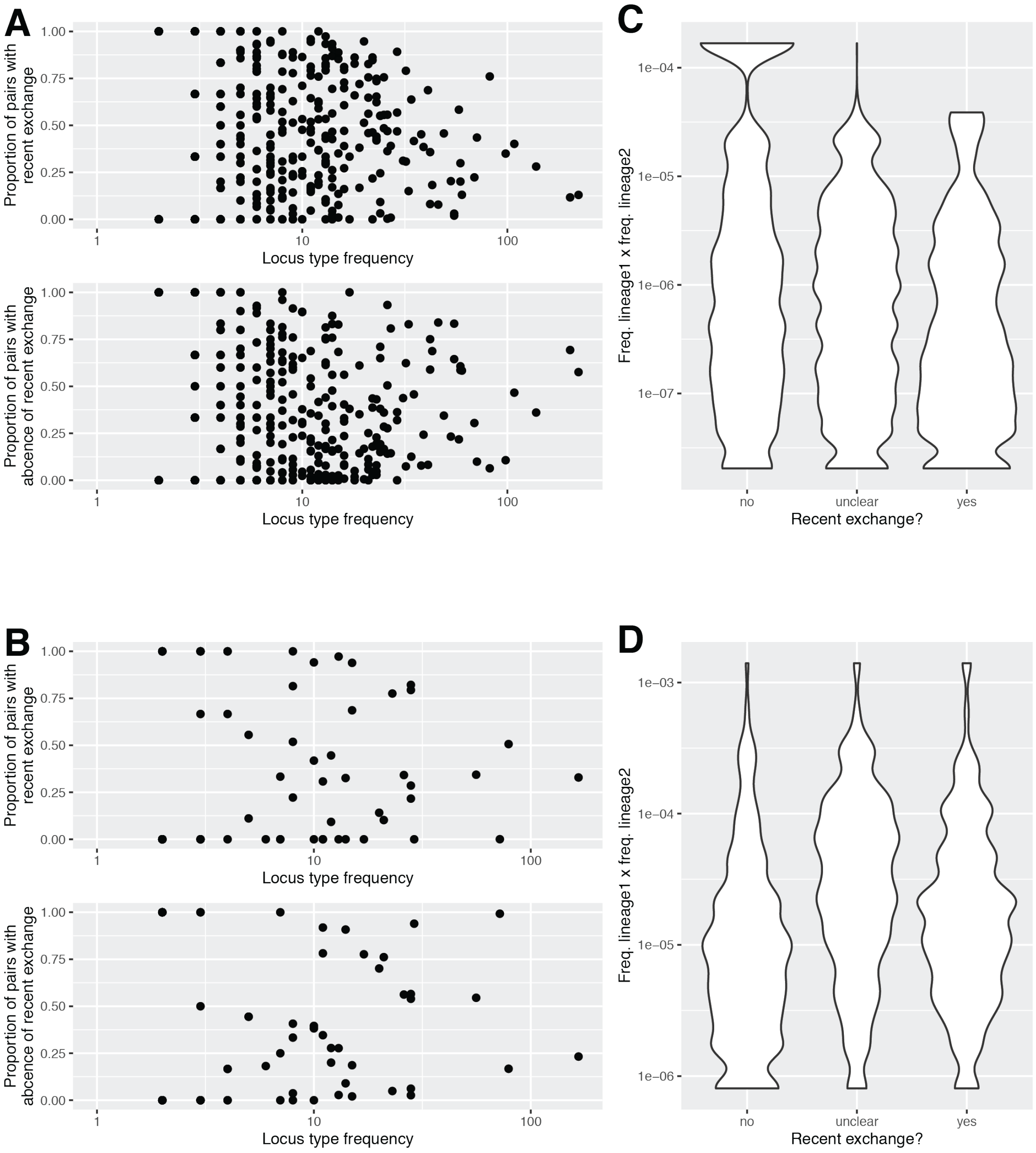


**Figure S12.** (A) Relationship between the frequency of all locus-types found in the *cps* locus region and a proportion of all pairs with that locus-type between which it was recently exchanged (top) or between which it was not recently exchanged (bottom). (B) Same as panel A but for the *kps* locus region. Correlation statistics for panels A and B are shown in Table S4. (C) Violin plots show the distribution of lineage frequencies $f_i f_j$ for all pairs $i$ and $j$ sharing a locus-type in the *cps* locus region and belonging to a given class: absence of recent exchange (left), presence of recent exchange (right) and unclear (middle). (D) Same as panel C but for *kps* locus region.

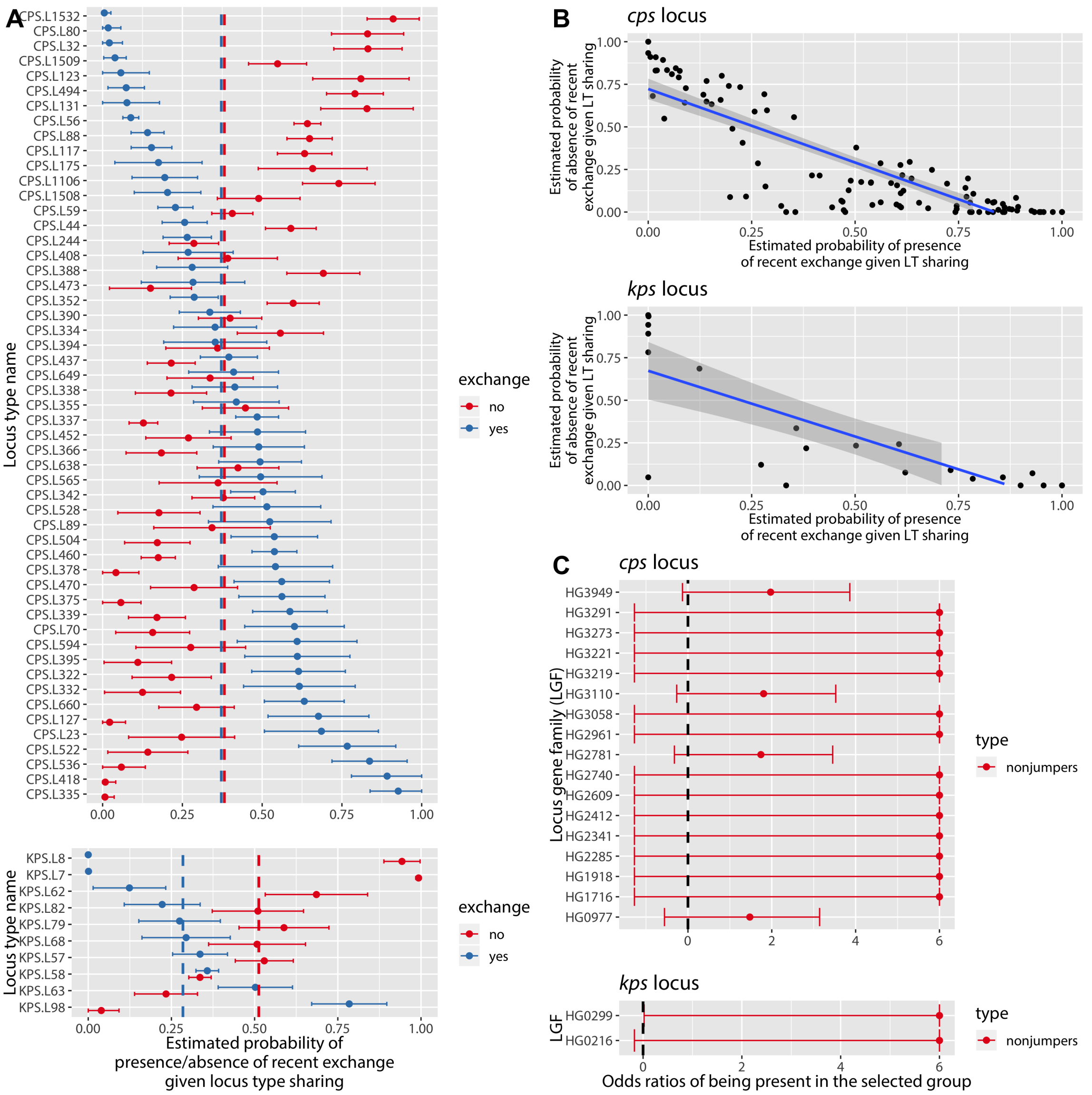


**Figure S13.** (A) Relative frequency of locus-type exchange via HGT (blue) and absence of exchange via HGT (red) for all representative pairs sharing that locus-type. Only locus-types shared between at least 100 pairs of $L_{0.004}$ lineages are shown. Bars show confidence intervals at the level of $1 - 0.05/n$, reflecting a Bonferroni correction due to multiple testing (*cps*: $n = 618$, *kps*: $n = 73$). Dashed line shows the estimated probability of locus-type sharing via recent exchange (blue) or in absence of recent exchange (red) for all lineage pairs. (B) Comparison of blue and red estimates from panel A but for all locus-types with a significant estimate. A linear model fit is shown. (C) Odds ratios for a given locus gene family (LGF) being associated with extreme jumpers (blue) and extreme non-jumpers (red). Blacked dashed line shows OR = 1. Bars show confidence intervals at the level of $1 - 0.05/n$, reflecting a Bonferroni correction due to multiple testing (*cps*: $n = 132$ for jumpers and $n = 58$ for non-jumpers, *kps*: $n = 40$ for jumpers and $n = 20$ for non-jumpers). Functional annotations and names of LGFs are shown in Table S5.

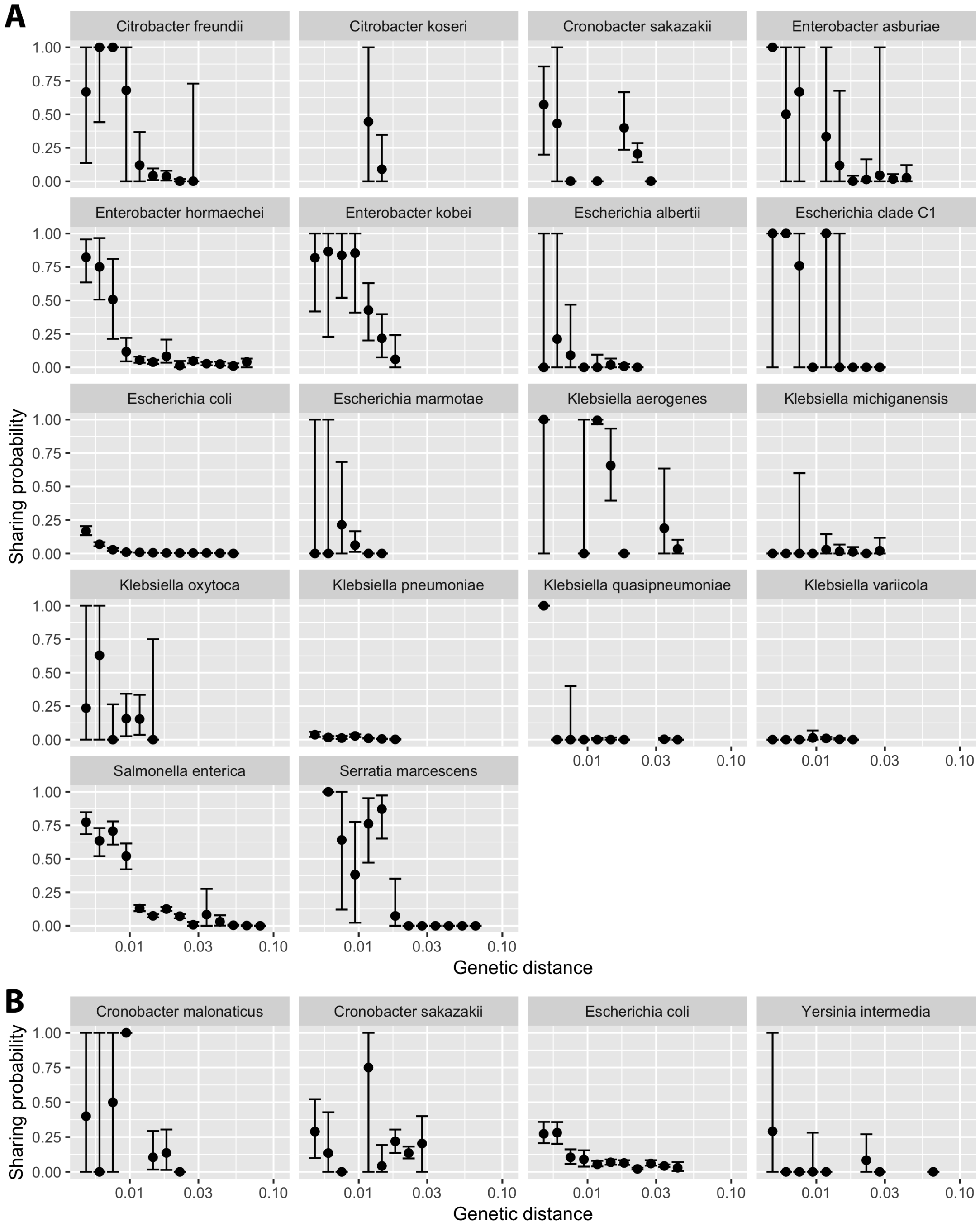


**Figure S14.** Estimated probability of within-species locus type sharing for each of the species as a function of genetic distance (1-ANI) between different lineages for (A) *cps* and (B) *kps*. Results show median (point) and 95% confidence intervals (error bars) of a bootstrap distribution obtained by sampling all isolates with replacement $n = 100$ times.

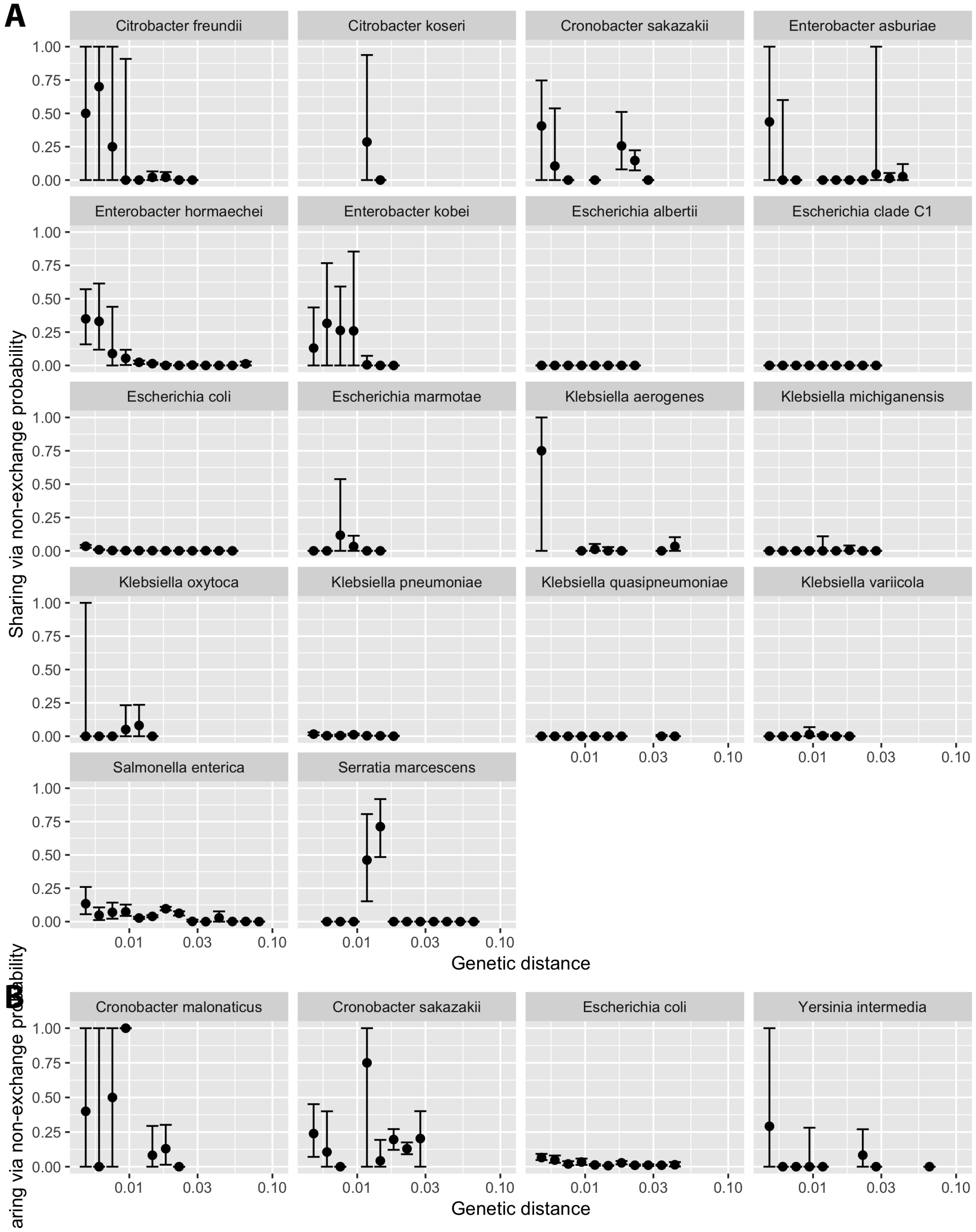


**Figure S15.** Estimated probability of within-species locus type sharing via no recent exchange (red category from Figure 3) for each of the species as a function of genetic distance (1-ANI) between different lineages for (A) *cps* and (B) *kps*. Plots are produced in the same way as in Figure S14.

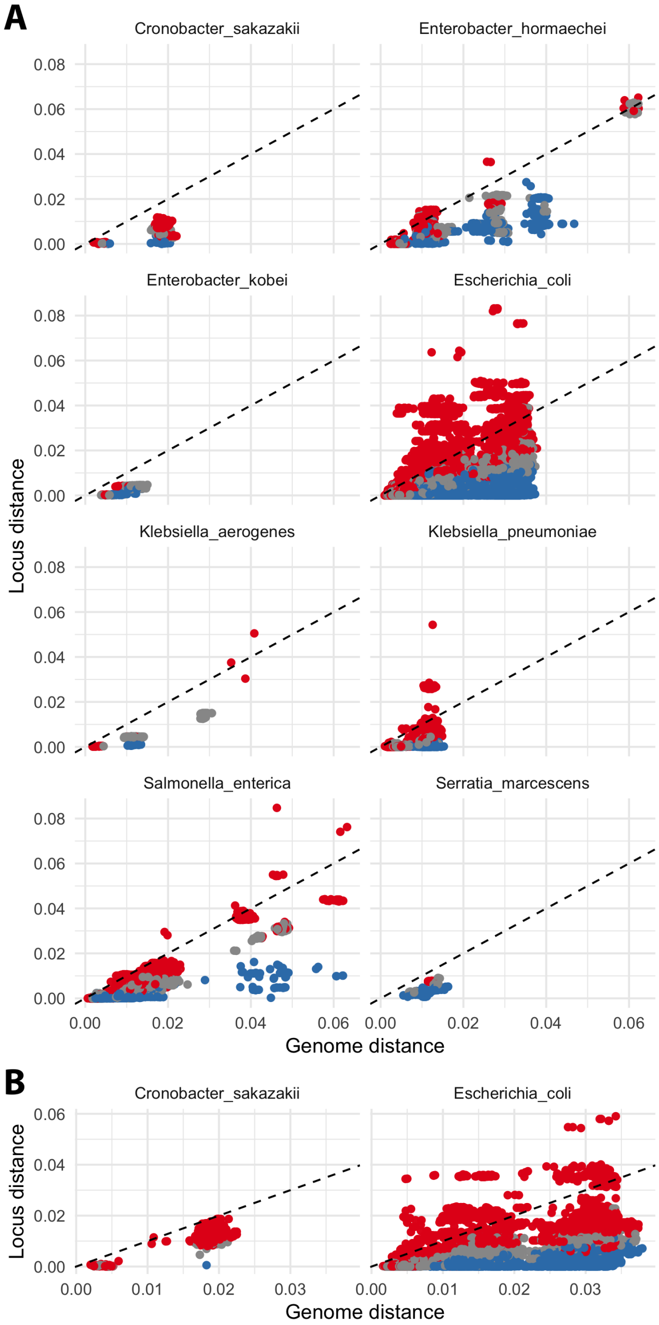


**Figure S16.** Each plot shows a relationship between the genome distance and the SP locus distance (1-ANI) in all within-species between-lineage pairs which share a *cps* locus (A) or *kps* locus type (B). Points are coloured according to the recombination test category, as indicated in Figure 3. The species shown correspond to those in Figure 5. Dashed line represents the $y = x$ curve.

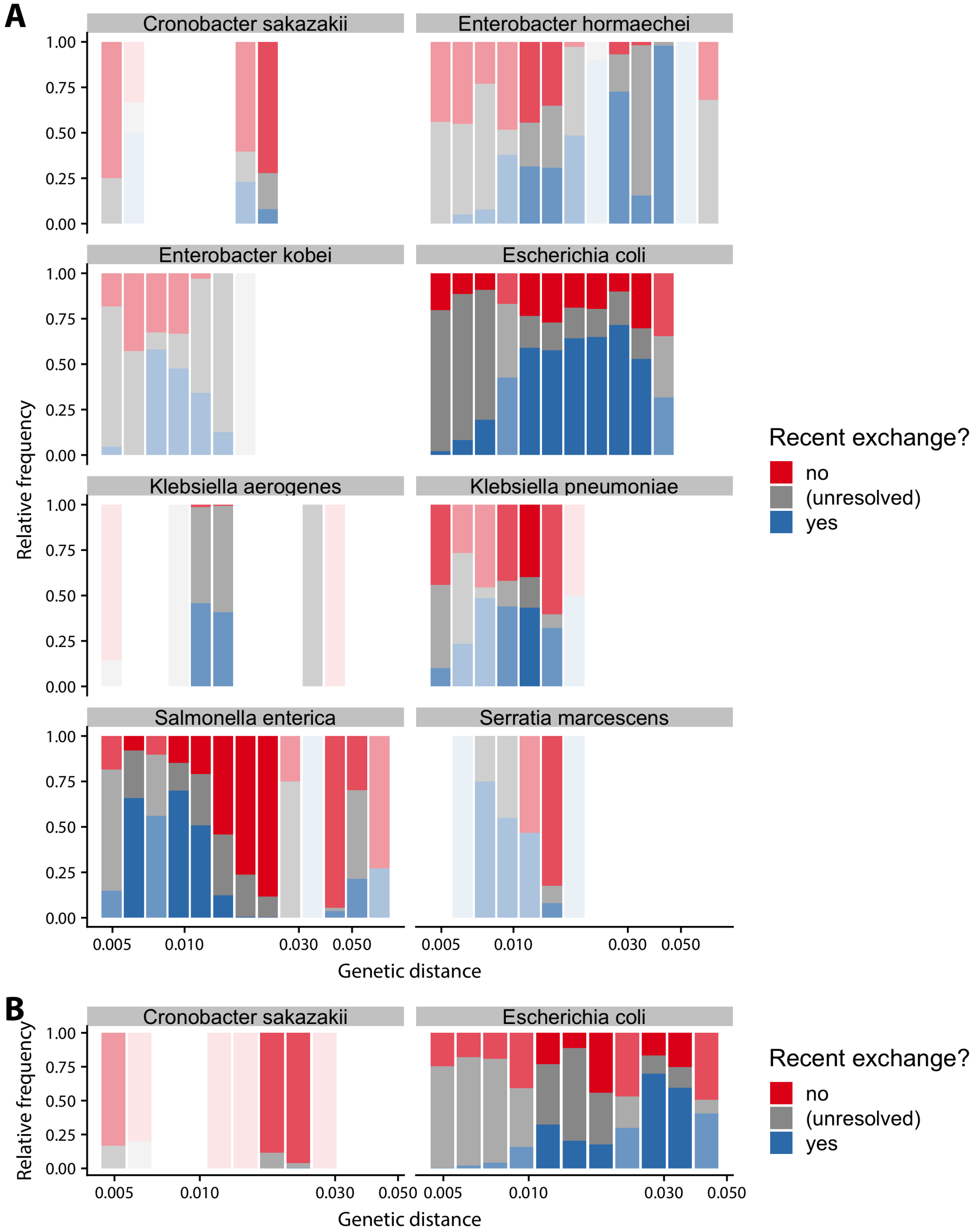


**Figure S17.** Relative frequency of the three categories in Figure S16 to locus-type sharing. Each bar represents a range of genome distance values binned into 15 categories, from 0.004 to 0.1. Transparency of each bar corresponds to the sample size $N$ (number of pairs) in that bin, with 10% opacity for $N \leqslant 10$, 40% opacity for $100 > N > 10$, 70% opacity for $1000 > N \geqslant 100$, and 100% opacity for $N \geqslant 1000$.

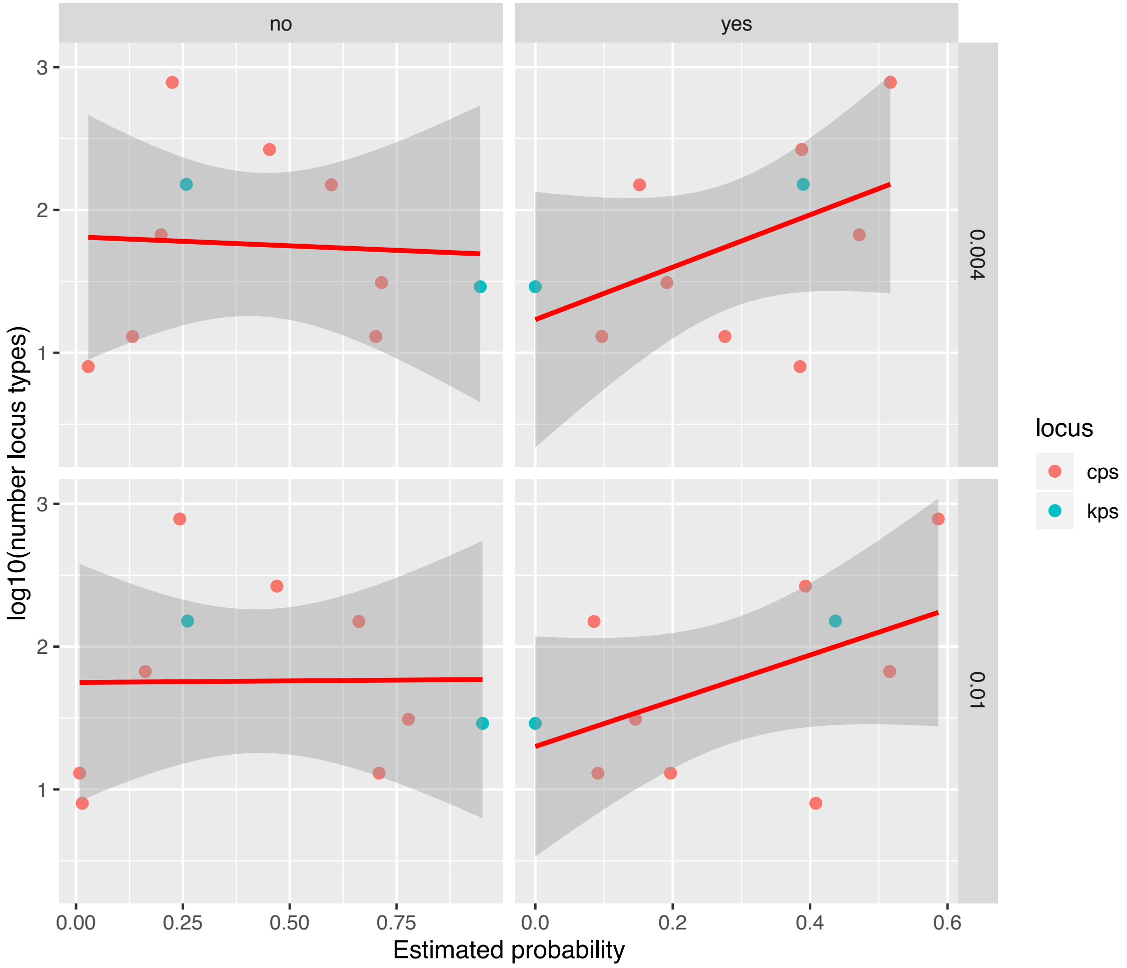


**Figure S18.** Correlation between log10 of number of locus-types and the estimated probability of locus-type sharing due to absence (left) and presence (right) of recent exchange via HGT, as shown in Figure 5C, and for the $L_{0.004}$ (top) and $L_{0.01}$ (bottom) lineage definition. No correlation was found for the absence of recent exchange variable; a borderline weakly positive correlation was found for the presence of recent exchange variable (Pearson correlation; $L_{0.004}$: $R^2 = 0.40$, $p = 0.050$; $L_{0.01}$: $R^2 = 0.41$, $p = 0.048$).

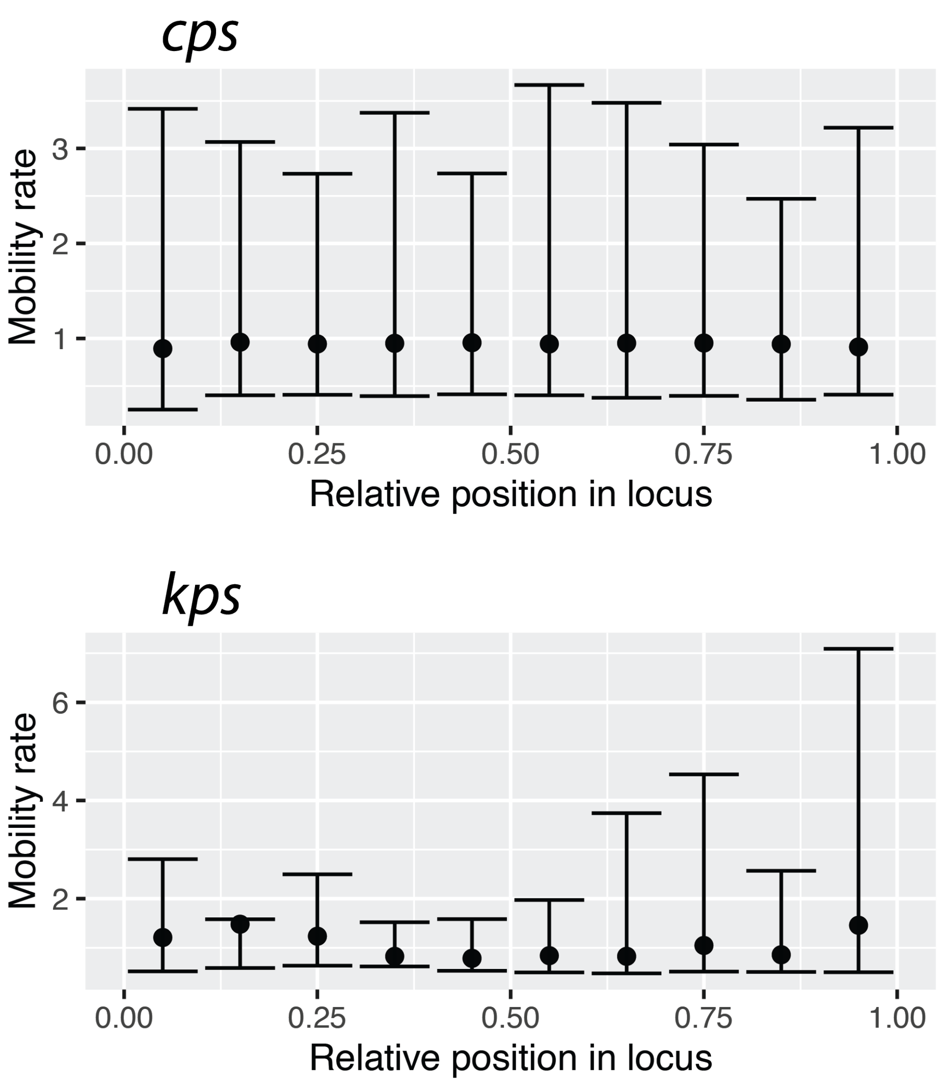


**Figure S19.** Each point shows the median gene mobility rate, as measured by GLOOME, for all LGFs assigned to a given category depending on their relative position. Position for each LGF in a given lineage was calculated as an average of all positions of that LGF in each isolate, divided by the total number of LGFs in that isolate.

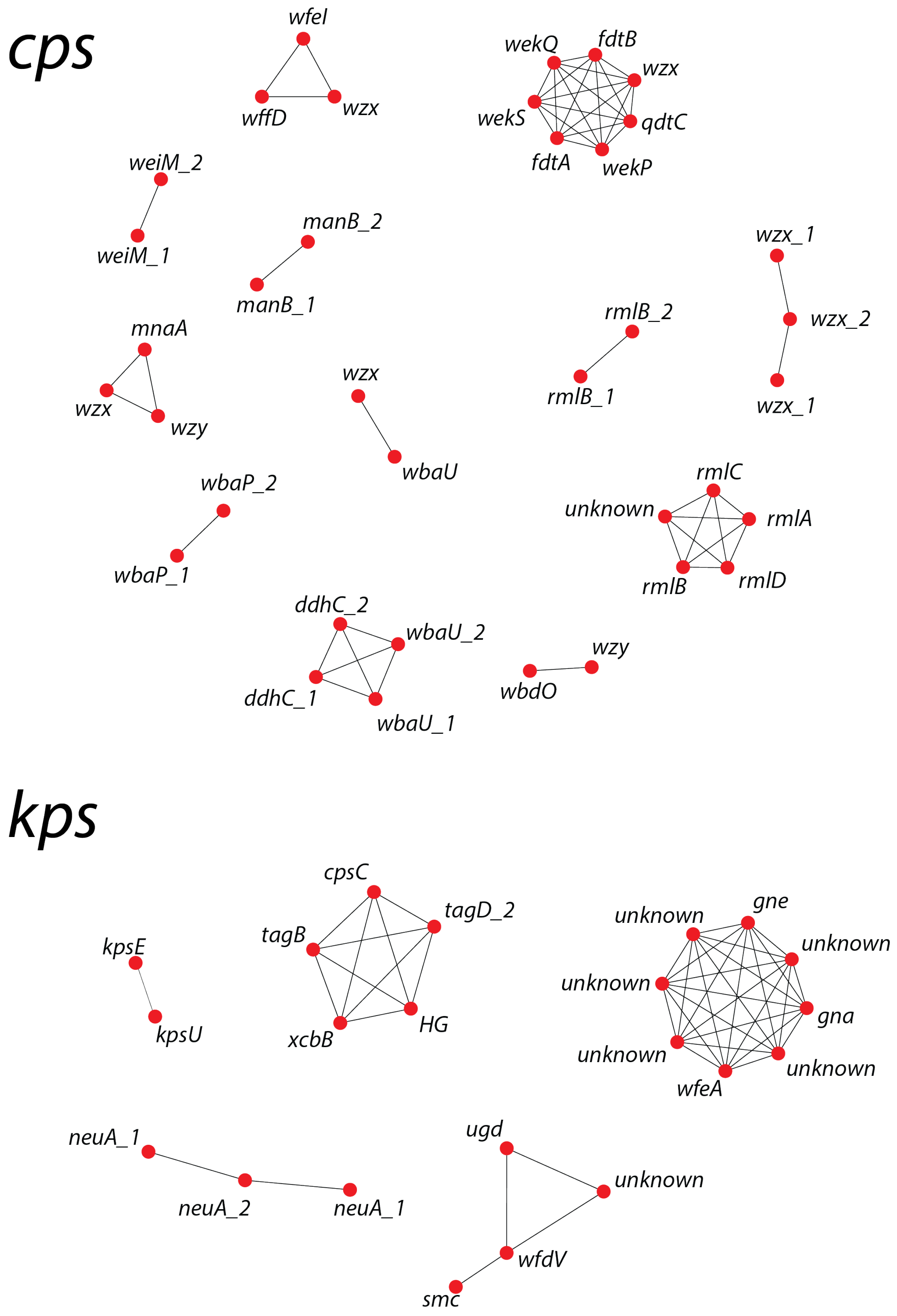


**Figure S20.** Network shows clusters of co-mobility between LGFs in the *cps* (top) and *kps* (bottom) region. Nodes show LGFs and edges link those LGFs pairs with the co-mobility coefficient significantly greater than 0.5. The most common gene names in a LGF are shown.

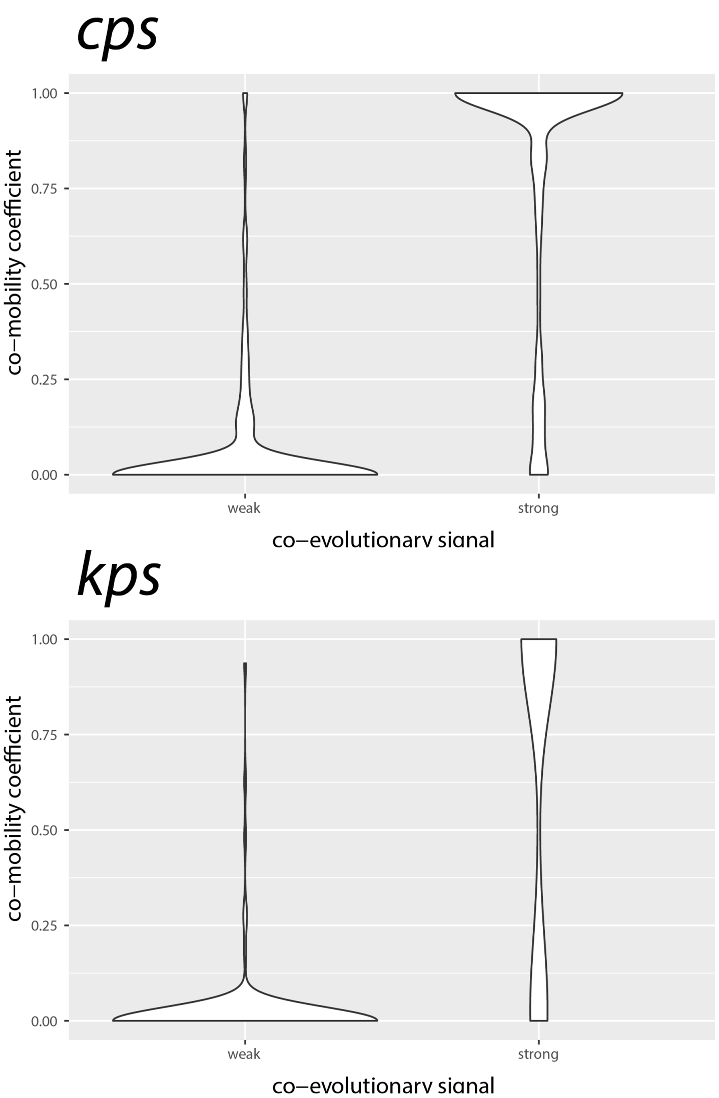


**Figure S21.** Violin plots of co-mobility values for LGF pairs with a weak ($< 0.2$, $n = 2305$) vs. strong ($\geqslant 0.8$, $n = 249$) co-evolutionary signal in the two locus regions. Only LGF pairs with a significantly positive value of the Mantel test coefficient were considered.

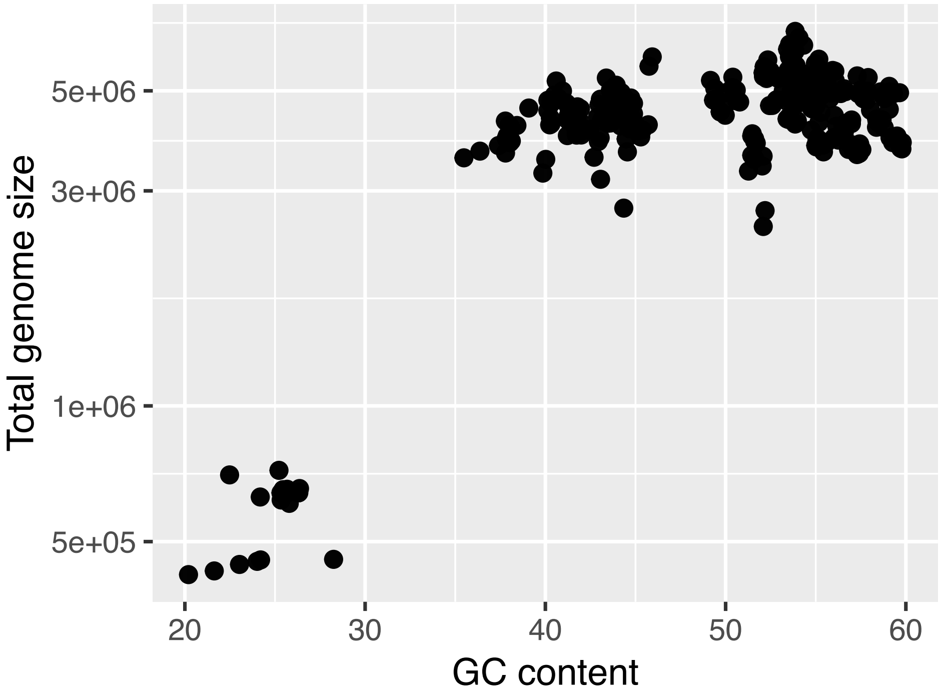


**Figure S22.** Distribution of the assembly GC content (X-axis) and total genome length (Y-axis) of all isolate clusters at the ANI distance of 0.15 with 20 isolates of less per cluster ($n = 361$). The bottom-left group of points belongs to the *Buchnera/Wigglesworthia* genus-group.

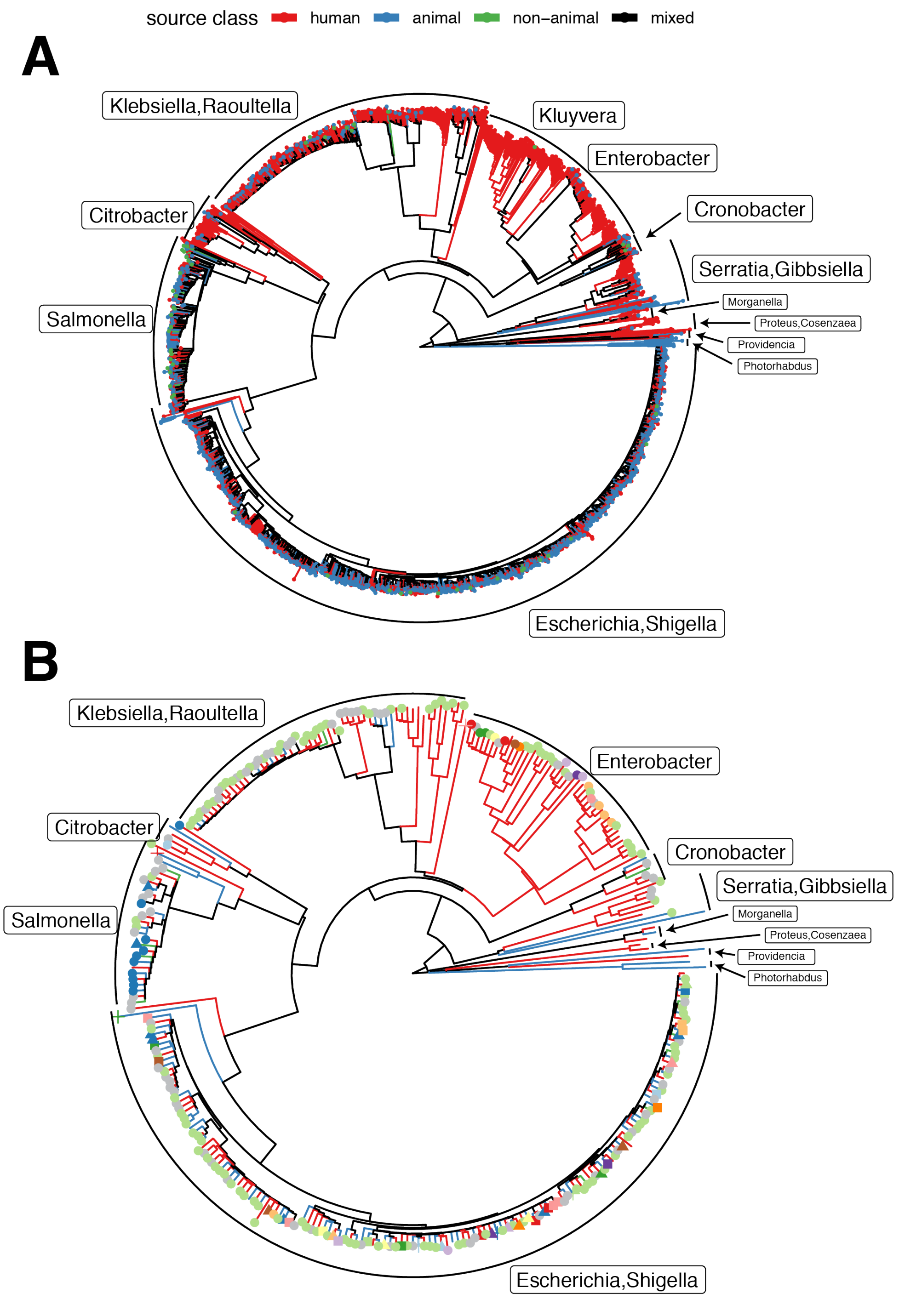


**Figure S23.** (A) Neighbour-joining tree of 1841 taxa across the Enterobacteriales order, coloured according to the assigned environmental class (human, animal or non-animal), as described in Methods. Taxa which have not been assigned to one of the classes have been omitted from the tree. Each branch of the tree is coloured in black unless all taxa downstream of a node to which that branch is leading belong to the same environmental class (with the same corresponding colour as the taxa). (B) A random subset of 300 taxa from the tree in panel A. A combination of the shape and colour next to each taxon represents a unique locus family (connected components in Figure 2).

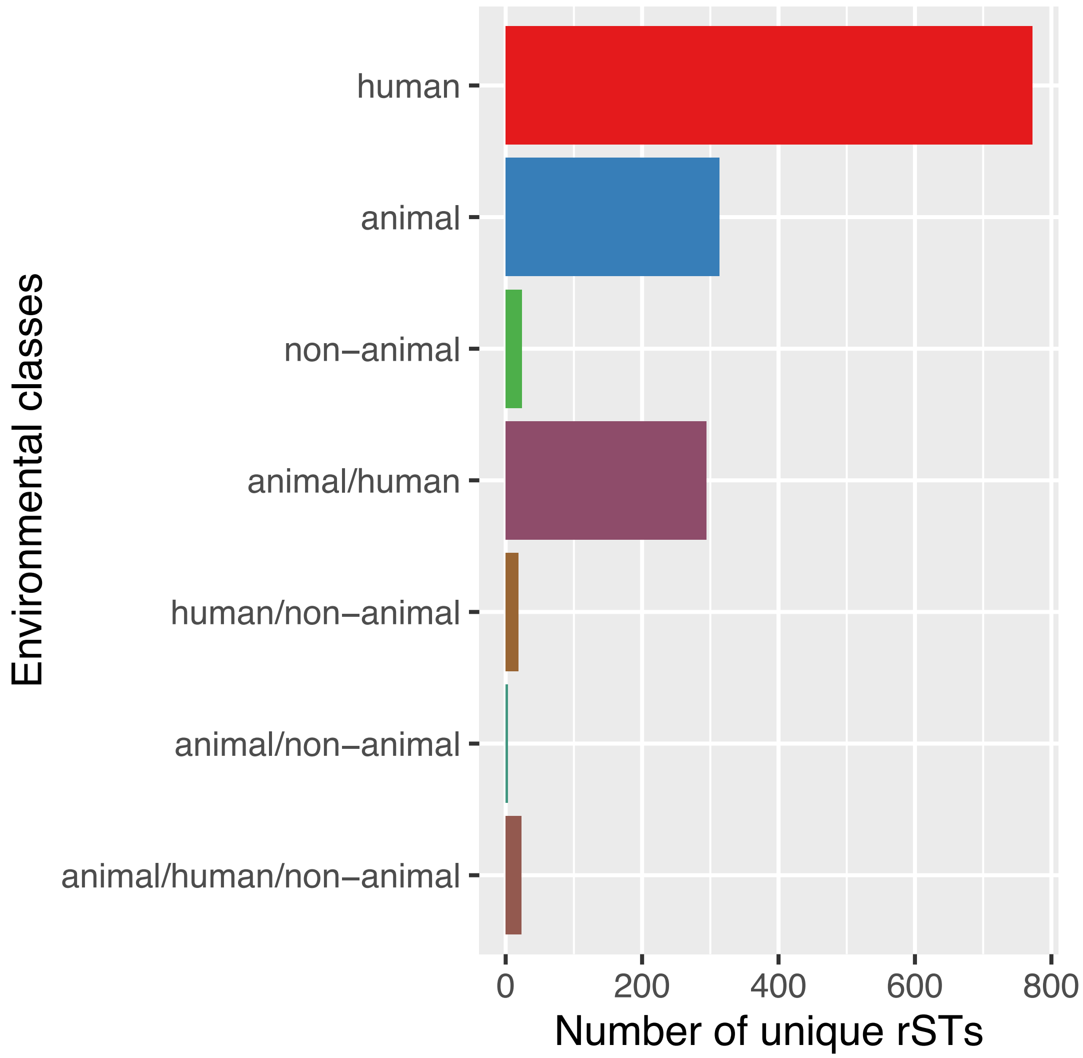


**Figure S24.** The bar plot shows the distribution of environmental classes across ribosomal sequence types (rSTs). Amongst 17,142 isolates which had an assigned environmental class, as described in Methods, there were 5,602 unique rSTs. Of those, 1,448 rSTs featured more than a single isolate, and these are the ones shown in the figure. Almost a quarter of those rSTs ($n = 339$, 23%) were assigned to multiple environmental classes.

**Table S1.** Metadata of the 27,334 genomes used in the study. The data are provided as an external XLSX file.

**Table S2.** Table showing the estimated species jump-rates for 43 LGFs in Fig. 6C and sequence names used to produce the homology group alignments. The sequence names can be found in the supporting datasets via Figshare. The data are provided as an external CSV file.

**Table S3.** Table showing assignment of 27,334 isolates to environmental classes, as described in Methods. The data are provided as an external XLSX file.

| | *cps* | *kps* |
|---|---|---|
| recent exchange $\sim$ frequency | $\rho = 0.071$, $p = 0.079$ | $\rho = 0.61$, $p = 0.060$ |
| absence of recent exchange $\sim$ frequency | $\rho = 0.21$, $p = 1.8 \times 10^{-7}$ | $\rho = 0.14$, $p = 0.23$ |

**Table S4.** Spearman correlation results between locus-type frequency in a representative dataset and a propensity of the locus-type to undergo recent exchange given sharing. Only unique combinations of locus-type and pairwise $L_{0.004}$ lineages were considered.

**Table S5.** List of all locus gene families (LGFs) present in extreme jumpers and extreme non-jumpers, together with the estimated odds ratios of association of these LGFs with jumpers/non-jumpers. The first tab lists all LGFs, and the second shows functional annotations obtained using `EggNOG-mapper` for LGFs with $p < 0.01$. The data are provided as an external XLSX file.
